# A recombinant expression system for the Variant Surface Glycoproteins enables structural elucidation of challenging classes of trypanosome proteins

**DOI:** 10.64898/2026.11.19.767849

**Authors:** Monique van Straaten, Johan Philip Zeelen, Kristin Degen, Julia Blesch, C. Erec Stebbins

## Abstract

The African Trypanosome (*Trypanosoma brucei*) persists in the mammalian host by periodically altering its surface coat with antigenically distinct versions of the **V**ariant **S**urface **G**lycoprotein (VSG). This process is called antigenic variation and produces repeated cycles of immune clearance and parasite VSG switching, leading to high morbidity and mortality in the absence of chemotherapeutic intervention. Elucidating how variant VSGs evade immune clearance is thus central to understanding trypanosomiasis. Progress in studying the large repertoire of thousands of VSG proteins has been hindered by the challenge of producing these coat proteins. Here we report the use of suspension HEK293 cells for this purpose, a system with robust yields that preserves many key elements of VSG structure and post-translational modifications, verified by expressing, purifying, and determining the crystal structures of the well-characterized proteins VSG2 and VSG3. We then applied this heterologous expression system to the production of several VSGs that had failed to express natively in *T. brucei,* including truncated VSG-Related Proteins (VRPs) from *T. brucei* and VSGs from *T. vivax*. This allowed for the determination of the structures of the *T. brucei* VRP VSG1619 as well as the VSG F9WSW0 from *Trypanosoma vivax (T. vivax).* The structure of VSG1619 shows similarities to the upper-lobe structures of B-class VSGs, but the monomers do not assemble into either the typical A-class dimer or B-class trimer in solution or in the crystals. The structure of *T. vivax* F9WSW0 is notable as a divergent member of the Class-A group that was developed from structures of *T. brucei* VSGs, with a canonical dimer but differences in fold, disulfide placement, and *N*-linked glycosylation. The described expression system provides additional insight into the family of VRPs and opens the door to general progress with the VSGs of more divergent trypanosome species through recombinant protein production.

**Author Summary:** The African trypanosome is a single-celled parasite causing many diseases in animals and sleeping sickness in humans. It can avoid the immune system indefinitely because it repeatedly changes the protein coat on its surface to a different variant that is unrecognized. There are thousands of these variants in the genome of the parasite. Understanding the molecular structure of these variants is critical to understanding how they avoid immunity. Making these variants for scientific study can be challenging, and we report here a new method using a very common protein expression system that successfully achieves this goal. We test it on well-studied variants showing that the system produces protein that closely matches the material produced in trypanosomes. We then use this system to determine the atomic structures of two variant proteins that have not been successfully produced before, one of them from a distantly related species of this parasite. This system provides a robust protein production pathway to examine the trypanosome surface proteins and should facilitate the broadening of scientific knowledge of these variants.

## INTRODUCTION

African trypanosomes (e.g., *Trypanosoma brucei*, including *T. b. gambiense* and *T. b. rhodesiense*, *T. congolense, T. vivax*) are single-celled, flagellated, eukaryotic parasites primarily found in sub-Saharan Africa that infect a wide range of mammals causing diseases such as *nagana* in livestock and sleeping sickness in humans [1,2]. They are spread by the bite of the tsetse fly vector (*Glossina* spp.) [3]. Like related trypanosomes such as *Trypanosoma evansi* (closely related to *T. brucei* and the causative agent of *surra* in camels, horses, and dogs), they rely on antigenic variation of their surface coat to evade the adaptive immune response [4–6]. Across the geographic regions impacted by various trypanosome species, the organisms cause significant morbidity and mortality, especially in animals, with a significant economic impact on the nations involved [7,8].

Trypanosomes live freely in the blood of the host organism and are therefore constantly exposed to the full surveillance and assault of the immune system [3]. The parasite escapes host adaptive immune clearance by altering its surface coat on a regular basis. The trypanosome surface is carpeted by roughly ten million copies of the Variant Surface Glycoprotein (VSG) in a continuous monolayer ∼12–15 nm thick. Accounting for approximately 90-95% of total surface protein, this dense coat is highly immunogenic, leading to robust neutralizing antibody responses [9,10]. However, the infection continues because some percentage of trypanosomes will switch to antigenically distinct VSG proteins from a genomic repertoire of thousands of such genes. The immune system is naïve to these new parasites with a novel coat, and this leads over time to recurring peaks and valleys of parasitemia as the immune system chases the changing antigenic landscape on the surface of the infecting organism [11].

Therefore, understanding the “epitope space” of the VSGnome (the genomic repertoire encoding Variant Surface Glycoproteins) is central to understanding the long-term immune evasion strategy of this organism. Previously, we have experimentally determined the structures of numerous VSGs from *T. b. brucei* and used deep-learning computational models such as AlphaFold [12,13] and ESMFold [14] to characterize and classify the broad genomic repertoire of the N-terminal domains (NTDs) of VSGs from this organism [15], the largest and most membrane-distal domain shown to be the most immunogenic [16–19]. However, there are “outlier” VSG and VSG-like sequences in the genome, many of which we have found challenging to express natively on the *T. brucei* surface. Additionally, VSGs from other species often show significant divergence from those in *T. brucei*, raising the possibility of structural and antigenic differences beyond those seen to date. VSGs from other trypanosome species that are heterologously cloned into *T. brucei* (one of the few culturable trypanosome species with generally robust yield) can be challenging to express [20]. Finally, protein modeling has often struggled with these more divergent VSGs, as evinced by many database entries with low confidence scores and unrealistic structures [15].

Previous work has shown that VSG NTDs can be expressed in heterologous eukaryotic expression systems. For example, *Pichia pastoris* was used to express RoTat1.2 from *Trypanosoma evansi* and the T. b. gambiense VSGs LiTat 1.3 and LiTat 1.5, the latter two also expressed in *Leishmania tarentolae* [21–23]. While these recombinant versions showed serological activity, these studies characterized the recombinant proteins only through indirect methods such as electrophoresis and antigenicity.

We have employed a heterologous expression system in Human Embryonic Kidney (HEK) cells that allows for the production of VSGs recalcitrant to production on the *T. brucei* surface. We established the functionality of this system by expressing, purifying, and determining the crystal structures of the well-studied VSG2 and VSG3 proteins, showing that these maintained their protein architectural characteristics and oligomeric assemblies, as well as many post-translational modifications and ligand binding properties (e.g., *N*-linked glycosylation, VSG2 binding to calcium, but not *O*-linked glycosylation in VSG3). We then used this system to express and determine the structures of a member of a class of truncated VRPs from *T. brucei* as well as the VSG F9WSW0 from *T. vivax*.

## METHODS

### Cloning

VSG N-terminal domains without the signal peptide (VSG2_27-391_, VSG3_19-393_, VSG1619_23-260_, VSG3327_33-267,_ henceforth denoted with “NTD”, e.g., VSG2-NTD) were amplified from plasmids containing the full-length proteins and cloned into an appropriate vector for producing proteins under the control of a CMV promoter and adding the mouse immunoglobulin H chain V-region signal peptide (MGWSCIILFLVATATGVHS; vector was kindly provided by Mirjana Lilic, S1 Fig, derived from VRC01[24]). NEBuilder HiFi Assembly master mix (#E2621) was used for cloning. The VSG3 and VSG1619 constructs contained a C-terminal hexahistidine sequence (His-Tag). VSG3327 mutants N50A and N192A were prepared by site-directed mutagenesis using QuikChange Lightning (Agilent Technologies)

For cloning the *T. vivax* proteins, DNA constructs were ordered from ThermoFisher GeneArt services. From the Uniprot database (uniprot.org), three Class-A/Fam23 (F9WKJ2/TvY486_0006500, F9WNN2/TvY486_0018500 and F9WSW0/TvY486_0035080) and three Class-B/Fam24 (F9WSQ1/TvY486_0034390, F9WW11/TvY486_0005140 and F9WV42/TvY486_0043480) VSGs were selected. The sequence containing the N-terminal domain was selected based on an analysis of signal peptide prediction (https://services.healthtech.dtu.dk/services/SignalP-6.0/). Using ESMFold and AlphaFold2/3, the end of the N-terminal domain was chosen through structural considerations, including pTM and ipTM scores comparing different oligomeric models. The cloning of the NTDs (F9WKJ2_28-363_, F9WNN2_18-365_, F9WSW0_23-369_, F9WSQ1_20-422_, F9WW11_21-334_ and F9WV42_21-363_) was performed as above (the number scheme may differ from the database, as protein start residues are not necessarily correctly annotated – see discussion in results with respect to F9WSW0, for example).

### Protein Expression and Purification

VSGs were produced in suspension (FreeStyle) 293-F cells by transfection using 293fectin (Thermo Fisher 12347-019). Cells were incubated at 37°C, 8% CO_2_ on an orbital shaker at 130 rpm for 5 days. Cells were removed by centrifugation at 1200 *g* for 10 minutes. Supernatants containing the protein were sterile filtered and concentrated in an Amicon stirred cell.

### Purification of VSG2-NTD

After buffer exchange to 10 mM sodium phosphate, pH 8, the sample was loaded onto a 15 ml Q-sepharose Fast-flow column (Cytiva), equilibrated and washed with 10 mM sodium phosphate pH 8. VSG2-NTD was eluted with 300 mM NaCl, 10 mM sodium phosphate, pH 8, concentrated and further purified by gel filtration (GE Healthcare, Superdex 200 Increase 10/300 GL column) in 10 mM HEPES, 150 mM NaCl, pH 8. Fractions containing the protein were concentrated in centrifugal units to about 2 mg/ml for crystallization. Samples were analyzed by SDS-PAGE.

### Purification of VSG3-NTD and VSG1619-NTD

After buffer exchange to binding buffer (300 mM NaCl, 50 mM sodium phosphate, 20 mM imidazole, pH 8), samples were incubated with Ni-NTA agarose (Qiagen) overnight at 4°C. Gravity flow columns were used to collect the Ni-NTA agarose, columns were washed with binding buffer, the VSG N-terminal domains were eluted with 50 mM sodium phosphate, 300 mM NaCl, 250 mM imidazole, pH 8, concentrated in a centrifugal unit and further purified by gel filtration (GE Healthcare, Superdex 200 Increase 10/300 GL column) in 10 mM HEPES, 150 mM NaCl, pH 8. Fractions containing VSG3-NTD were concentrated in centrifugal units to 1.7 mg/ml for crystallization. VSG1619-NTD was concentrated to 4.9 mg/ml. Samples were analyzed by SDS-PAGE.

### Purification of *T. vivax* VSGs

Following purification, F9WSQ1, F9WSW0 and F9WW11 showed a clear band on the SDS-PAGE gel. For the purification the supernatant containing the protein was concentrated and diluted with a buffer containing 20 mM Tris/HCl 0.5 M NaCl, 1 mM MnCl_2_ and 1 mM CaCl_2_, pH 7.4 and further concentrated. The protein was purified on a 1 ml HiTrap Con A 4B column (Cytiva) eluted with 300 mM methyl-α-D-glucoside according to the manufacturer’s recommendation. The eluted protein was concentrated and further purified by gel filtration (GE Healthcare, Superdex 200 Increase 10/300 GL column) in 10 mM HEPES, 150 mM NaCl, pH 8. The fractions containing the protein were pooled and concentrated.

### Crystallization

VSG2-NTD was crystallized at 22°C by vapor diffusion using hanging drops formed from mixing a 1:1 volume ratio of the protein with an equilibration buffer consisting of 22-24% (w/v) PEG 1500, 100 mM Tris-HCl pH 8.5. For data collection, crystals were soaked in the same buffer augmented to 20% ethylene glycol, flash-cooled in liquid nitrogen.

VSG3-NTD was crystallized at 22°C by vapor diffusion using hanging drops formed from mixing a 1:1 volume ratio of the protein with an equilibration buffer consisting of 2 M ammonium sulfate, 100 mM ADA pH 6.5, 0.1 M MgSO_4_. For data collection, crystals were soaked in the same buffer augmented to 20% ethylene glycol, flash-cooled in liquid nitrogen.

VSG1619-NTD was crystallized at 4°C by vapor diffusion using hanging drops formed from mixing a 1:1 volume ratio of the protein with an equilibration buffer consisting of 17% (w/v) PEG 6000, 100 mM sodium citrate pH 5.5, 3.5% t-butanol. For data collection, crystals were soaked in the same buffer augmented to 25% ethylene glycol, flash-cooled in liquid nitrogen.

F9WSW0 could be crystallized by sitting drop, 0.62 µl protein (1.2 mg/ml) and 0.62 µl 25% PEG1500 and 100 mM MIB (malonic acid / imidazole / boric acid). pH 7.0. For data collection, the same buffer with 15% glycerol was used and the crystals were flash-cooled and stored in liquid nitrogen.

### Structure Determination

Native datasets of VSG2-NTD and VSG3-NTD were collected at the BESSY II synchrotron. The crystals diffracted to 1.75 Å (VSG2-NTD) and 1.66 Å (VSG3-NTD) and the structures were solved using the PHENIX software package [25,26] with PHASER [27] using PDB IDs 7P56 and 6ELC as search models for VSG2 and VSG3, respectively. The model was improved with multiple rounds of refinement in PHENIX-REFINE and manual model building in COOT [28]. For VSG1619 a native dataset was collected at the BESSY II synchrotron diffracting to 1.47 Å. The structure was solved using an AlphaFold3 model with all the loops removed as search model in PHENIX-PHASER and the model rebuilt using PHENIX-AUTOBUILD, followed by multiple rounds of refinement in PHENIX-REFINE and manual building in COOT. A dataset for F9WSW0 diffracting to 1.74 Å was collected at BESSY II and an AlphaFold3 model was used for molecular replacement. Using ModelCraft from the CCP4 package [29] the protein was rebuilt as the loops from the top domain were not predicted correctly. Several rounds of optimization using PHENIX-REFINE, COOT and PDB-REDO [30] were used to finalize the model.

### Data Availability

Coordinates and structure factors have been uploaded to the RCSB PDB (www.rcsb.org) with the following PDB IDs: 35EL for F9WSW0, 50OP for VSG1619, 50OU for VSG3, and 50PF for VSG2.

## RESULTS

### HEK-Based VSG Production and Purification

Based on a common transient-transfection plasmid system (Methods, S1 Fig) employed for many proteins such as antibodies [31], Human Embryonic Kidney (HEK) FreeStyle^™^ 293-F cells (Thermo Fisher Scientific) were used to express the NTDs of VSG2 and VSG3 as proof-of-principle variants from *T. brucei brucei* Lister 427 strain (Methods). Both proteins were successfully secreted into the media and purified either via affinity resins or other means (Methods, S1 Table, and for all the constructs described in this paper, S2 Fig, A-E). Expression of VSG2 was at much higher levels than VSG3, indicating protein-to-protein variability in yields, a phenomenon observed in the expression of antibodies and other proteins in such systems [32].

Size exclusion chromatography (SEC, S2 Fig, B and C) showed elution peaks for the N-terminal domain (NTD) similar to native preparations of VSG2 and VSG3. As expected, VSG2 eluted as a dimer and VSG3 as a monomer. Prior to further work (e.g., crystallography), we verified the presence of *N*-linked glycans on VSG2 by PNGase treatment (S2 Fig, panel E), which showed a drop in molecular weight consistent with carbohydrate linkages expected from proteins purified endogenously from *T. brucei*.

Therefore, from standard biochemical assays used to assess the isolation of these two VSGs, the HEK-produced material was comparable to that produced from trypanosomes themselves. However, these methods cannot ascertain if the folded structure is unperturbed, or whether the glycosylation is properly maintained in the new expression system. We thus undertook high-resolution structural studies of these heterologously produced VSGs to address these issues.

### Heterologously Expressed VSG Structures

The NTD of HEK-expressed VSG2 was crystallized and the 1.75 Å structure determined experimentally by X-ray crystallography (Methods, S2 Table, S2 Fig, A and B). By all assessments, the HEK cell–produced protein is very similar in structure, glycosylation, and ligand binding to that natively produced (Figure 1A). VSG2 adopts the A2-class dimeric oligomerization, consistent with SEC data, and the core architectural, 3-helix bundle fold characteristic of the VSGs, the upper and lower lobes, and disulfide bond positioning are all conserved. The HEK and *T. brucei* produced VSG2 structures align with a root-mean-square deviation (RMSD) over 2596 atoms of 0.29 Å. The *N*-linked glycan is positioned correctly, attached to Asn 289 of the protein in the lower lobe, and for the residues visible, identical to that seen in *T. brucei* produced material. Therefore, the exogenous system yields VSG2-NTD that is at atomic resolution basically indistinguishable from trypanosome-produced material.

**Fig. 1.**
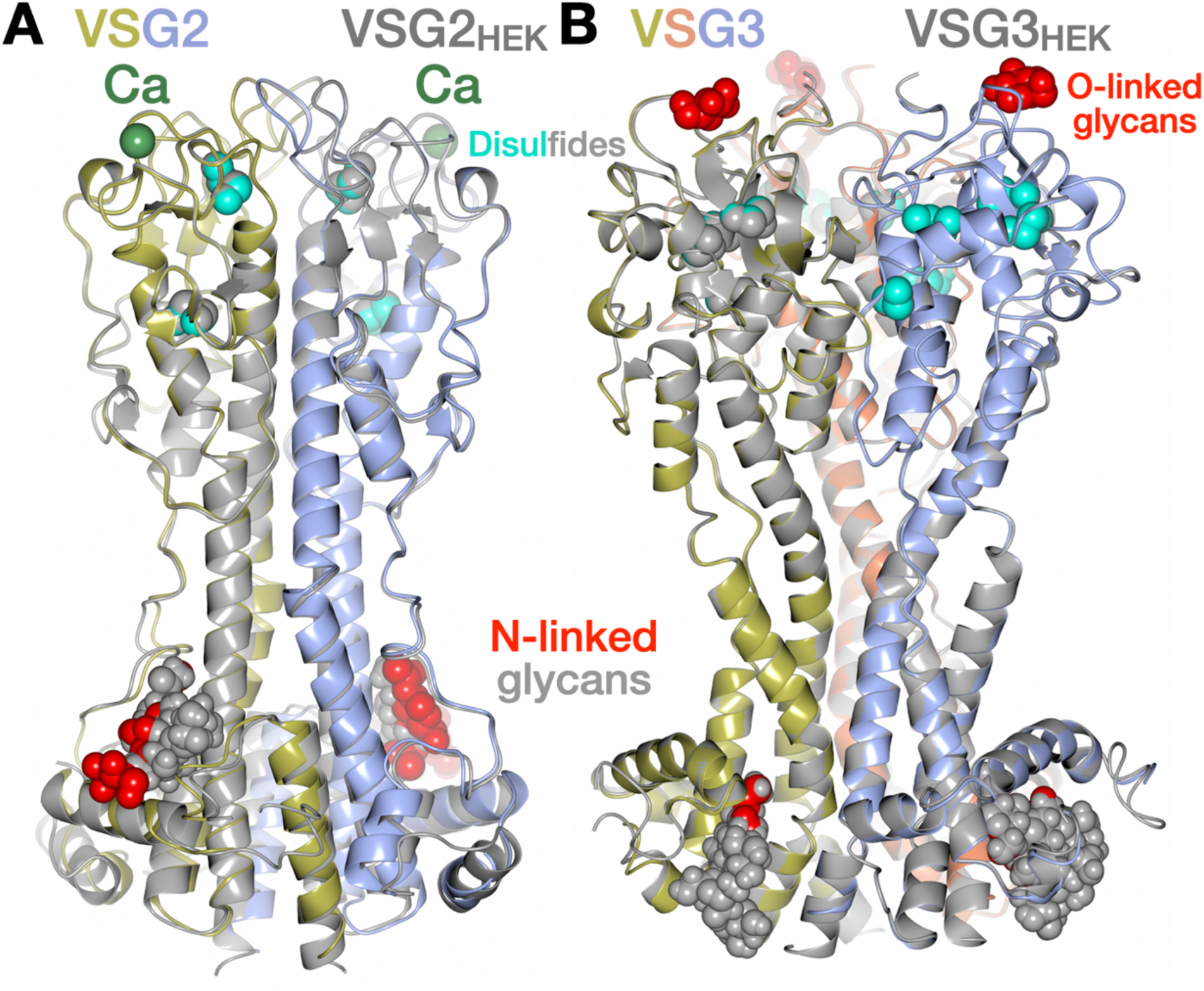
Comparing Native and Heterologously Produced VSG2 and VSG3. Superposition of VSG2 **(A)** and VSG3 **(B)** produced natively in *T. brucei* and in HEK-cells (VSG2_HEK_/ VSG3_HEK_). Native VSG2 is shown as a ribbon diagram in color (blue and gold for the different chains) and native VSG3 with blue, cyan and orange for the different chains, with a red, space-filling representation for the *N*-linked glycan in the lower lobe, disulfides in cyan, and the bound calcium in green. VSG2/3 _HEK_ are gray ribbon diagrams with gray space-filling atoms (note that the gray cannot be seen for the VSG2_HEK_-bound calcium ion as it overlaps perfectly with the ion in the trypanosome-produced material). Note that *O*-linked glycans in VSG3 produced natively in *T. brucei* are shown in red space-filling representation, but are not present in the structure of VSG3_HEK_.

Similar results were obtained for VSG3 with an exception with respect to *O*-linked glycosylation at the top of the protein. The NTD of HEK-expressed VSG3 was purified and crystallized, the 1.66 Å resolution structure determined experimentally by X-ray crystallography (Methods, S2 Table, and S2 Fig, A and C). While monomeric in solution (S2 Fig, panel C), and also in the asymmetric unit, the protein crystallized as a trimer on the three-fold axis of symmetry of the crystal, producing the trimer observed in all *T. brucei* class-B VSGs to date (the crystallographic packing in the space group I2_1_3 mirroring that seen in trypanosome produced VSG3, Figure 1B). The *N*-linked glycan is located correctly on Asn 67, the disulfides in the proper places, and the overall fold identical to endogenously produced protein with the RMSD between the structures produced by the two systems 0.25 Å over 365 amino acids of the trimer. A key difference between the endogenously and heterologously expressed VSG3 is the absence of any *O*-linked glycosylation in the latter. This is true for both *O*-linked residues seen in *T. brucei* produced protein (Ser 317 and Ser 319). The *O*-linked sugars of VSG3 have been shown to be immunodominant modifications that strongly modulate the host immune response to the pathogen [33,34]. In addition, only the B-class VSGs like VSG3 have been shown to have this modification, not the A-class proteins. A glycosyltransferase unique to *T. brucei* was suggested to target VSG3 with this modification [35], and therefore, the HEK-cell expression system that lacks this trypanosome enzyme might not be capable of making this modification (in contrast to the more broadly conserved *N*-linked glycosylation systems). However, it is also possible that the sugars were indeed added but not observed due to the labile nature of the modification, as has been seen in several VSGs that often lose this modification in purification or crystallization [33]. Therefore, while not definitive, it is likely that the HEK expression system does have a limitation in the area of trypanosome-specific sugar modifications that needs to be considered in its usage (which could perhaps be remedied through co-transfection of required glycosyltransferases).

### Heterologously Expressed Structure of Truncated VRP: VSG1619

The VSGnome is a complicated product of host-pathogen competition with genetic recombination and mutation occurring on both sides: on the one hand with antibodies in the adaptive immune system of hosts, and on the other with mosaic VSG formation by gene conversion in the trypanosome [11,16]. The products of these forces over millions of years have left large genomic repertoires of VSGs with proven functional coat proteins, truncated versions, VSGs with insertions, internal stop-codons, removal of signal sequences, and other often presumed debilitating mutations or alterations [36–38]. Although our previous work has characterized the VSGnome into a set of structural classes based on fold, topology, oligomerization, disulfide placement, and posttranslational modifications [15], a small percentage of VSG-like proteins in the *T. brucei* Lister 427 genome possesses minor or major alterations, putting them in various “outlier” subclasses.

One subset of VSG-related proteins (VRPs) is comprised of N-terminal domains smaller in size compared to standard VSGs yet showing detectable sequence similarity (S3 Fig, S1 File “ClustalAlignment_VRPsmall.pdf” and S2 File “VRP_truncated.xlsx”)[38]. While there has been some effort to link VRP size to function [39], it is still unclear how these truncated VRPs function in the trypanosome. Of those listed in S3 Fig, several properties show that they possess characteristics of functional membrane proteins (VRP_truncated.xlsx). Most score above predictive cutoffs for a consensus signal sequence and GPI anchor and do not harbor stop codons or obvious deleterious elements. AlphaFold3 predictions of structures build intact, VSG-like folds from these sequences (the bottom lobe missing, a truncated 3-helix bundle present, S3 Fig). While scattered across the genome, these shorter VSGs appear to be a closely related family generated from a common progenitor (S3 Fig, panel B). They possess nearly 50% sequence identity amongst each other, much higher than the average identity scores between VSGs. It is known that trypanosome viability is sensitively tied to VSG load on the surface [18,40], and should these be *bona fide* coat-VSGs used in antigenic variation, and not merely VRPs with other function(s), the difference in size between these truncated proteins and the more plentiful larger VSGs could require differential regulation of protein expression and/or trafficking. Overall, from these properties, it could be hypothesized that these VRPs might be expressed during infection.

Our own efforts have proven unsuccessful in cloning these VRPs into expression sites to bring these variants to the trypanosome coat. While certain aspects of how we do these expressions could explain the lack of success (monomorphic cells, using cassettes that are known to work for larger VSGs which possess different 3’ UTRs, etc.), a simple hypothesis could be that these are not surface VSGs but VRPs with other functions, and create non-viable trypanosomes when overexpressed from the Bloodstream Expression Sites (BESs). However, using the HEK system, we were able to examine whether stable protein can be made from these sequences, and if so, conduct biochemical and structural analyses of the expressed polypeptides.

Using the HEK system described above, we cloned, expressed, and purified VSG1619 (GenBank: AGH60539.1) to homogeneity (S1 Table, S2 Fig, A and D). The protein was soluble, stable, and amenable to crystallization. The structure was solved by X-ray crystallography to 1.47 Å resolution (Methods, S2 Table, Figure 2). While the AlphaFold3 model of VSG1619 was of moderate confidence in the 3-helix bundle region, the confidence score was low outside of this scaffolding core (S4 Fig, panel A) and efforts to solve the crystal structure by molecular replacement using the AlphaFold3 model failed. Trimming away the outer, low confidence regions of the structure that were not associated with the bundle fold led to models that were successful in molecular replacement (Methods), and the remainder of the structure was built through a combination of automated and manual methods.

**Fig. 2.**
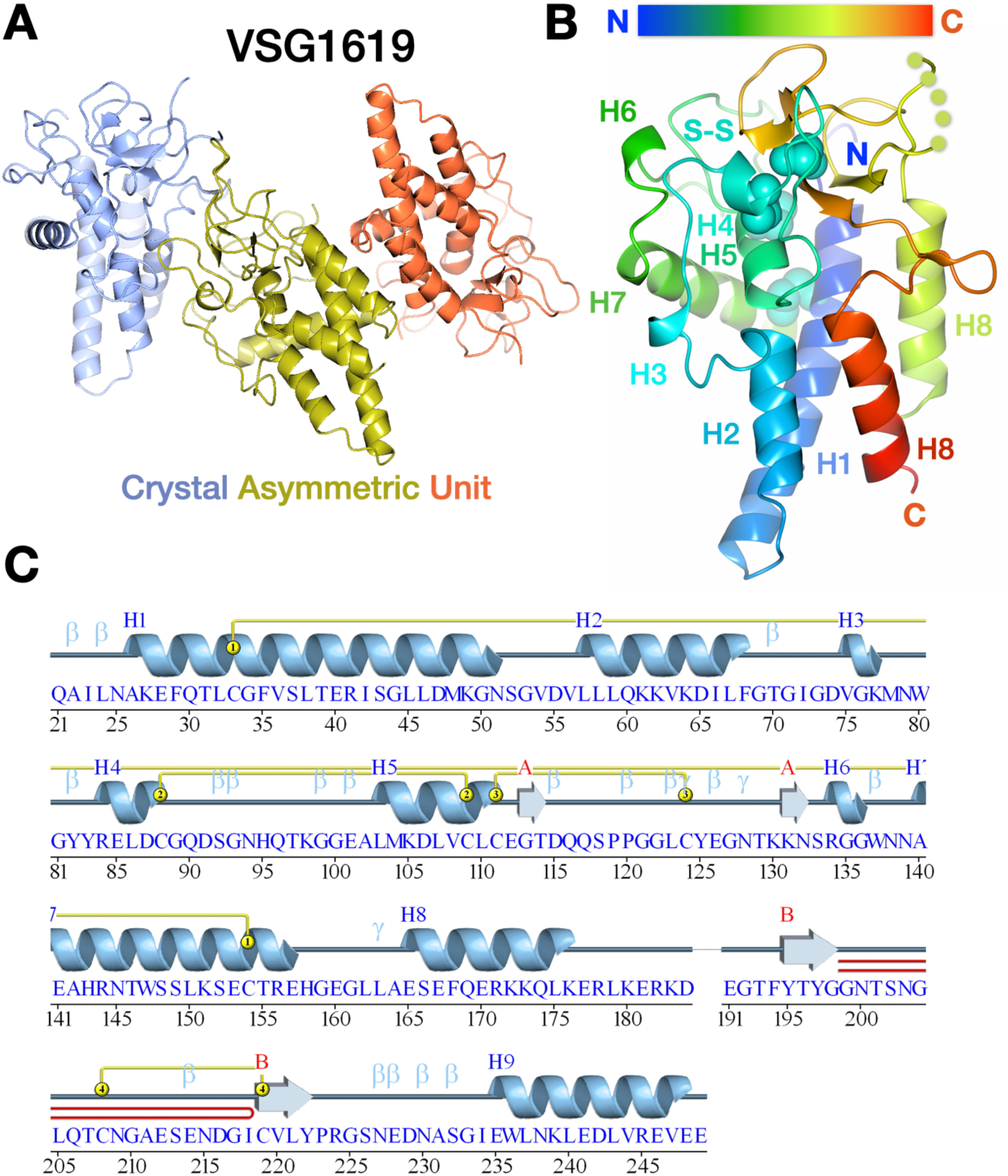
Structure of VRP VSG1619. (A) The contents of the crystal asymmetric unit are shown, each chain of VSG1619 shown as a ribbon diagram and given a different color. (B) Monomer of VSG1619 shown as a ribbon diagram presented with a color gradient from blue to red from the N-terminus (N) of the structure to the C-terminus (C). Disulfide bonds are shown as space-filling atoms in cyan and the helical secondary structural elements are labeled in accordance with panel. The five yellow circles indicate disordered residues for which no visible electron density existed. (C) showing a primary sequence-based view with protein secondary structure illustrated above. H and B refer to helical and beta-sheet geometry, respectively. “β” and “γ” indicate β-turns and γ-turns, respectively. Yellow lines above the sequence link bonded cysteines in disulfides. Red lines indicate β-hairpin turns.

The VSG1619 N-terminal domain (NTD) represents not only a sequence-truncated VRP, but also a structurally shortened member of the VSG super-family, divergent in several biochemical elements (Figure 2, A-C). To begin, there are three independent chains in the asymmetric unit of the crystal (Figure 2A) that do not assemble into any oligomeric arrangement that mirrors known full-length VSGs, either the dimeric arrangement of A-class VSGs or the trimeric arrangement observed in B-class VSGs, where monomers interact along the long-axis of the 3-helix bundle. Such alignment geometry is absent in the asymmetric unit of VSG1619 crystals. In addition, no dimeric or trimeric form of A- or B-class is present in the crystal packing through crystallographic symmetry operators. Coupled to the monomeric biochemical behavior in the purification of VSG1619 (S2 Fig panel D), these data suggest that these VRPs are monomeric as a subclass. Should they be expressed on the surface as VSGs, the unusually high densities and two-dimensional constraints of membrane packing could lead to different associative patterns, although, the AlphaFold confidence scores (ipTM) for oligomers are lower by significant margins than for assemblies of known Class A and Class B VSGs (S4 Fig, B and C). Alignments between the individual monomers in the crystal structure are around 0.5 Å over the Cα atoms, whereas the superposition with the AlphaFold 3 model is nearly 2 Å, with most of the divergence outside the three-helix bundle (S4 Fig, panel D).

Each chain in the asymmetric unit comprises ∼221–227 ordered residues and adopts a compact, single-lobe architecture (Figure 2B). The longest molecular dimension is ∼55-60 Å, with the fold organized around a mixed α/β scaffold capped by solvent-exposed loops that possesses a shortened extension of the 3-helix bundle of the VSG family. The topology best corresponds to the “upper lobe” defined for full-length B-class VSGs, while features contributed by the canonical “lower lobe” are absent in VSG1619. Other conserved B-class features include the location of disulfide bridges as well as the alignment of the fold and position in a structural dendrogram [41] (Figure 3 and S3 Fig, panel B). Therefore, while the absence of a lower lobe hampers the topological assignment of VSG1619 to either the A- or B-classes, the strongly conserved B-class elements still present in the remaining protein support the hypothesis that these truncated VSGs can be viewed as a divergent subclass of B-class VSGs (or VRPs more generally).

**Fig. 3.**
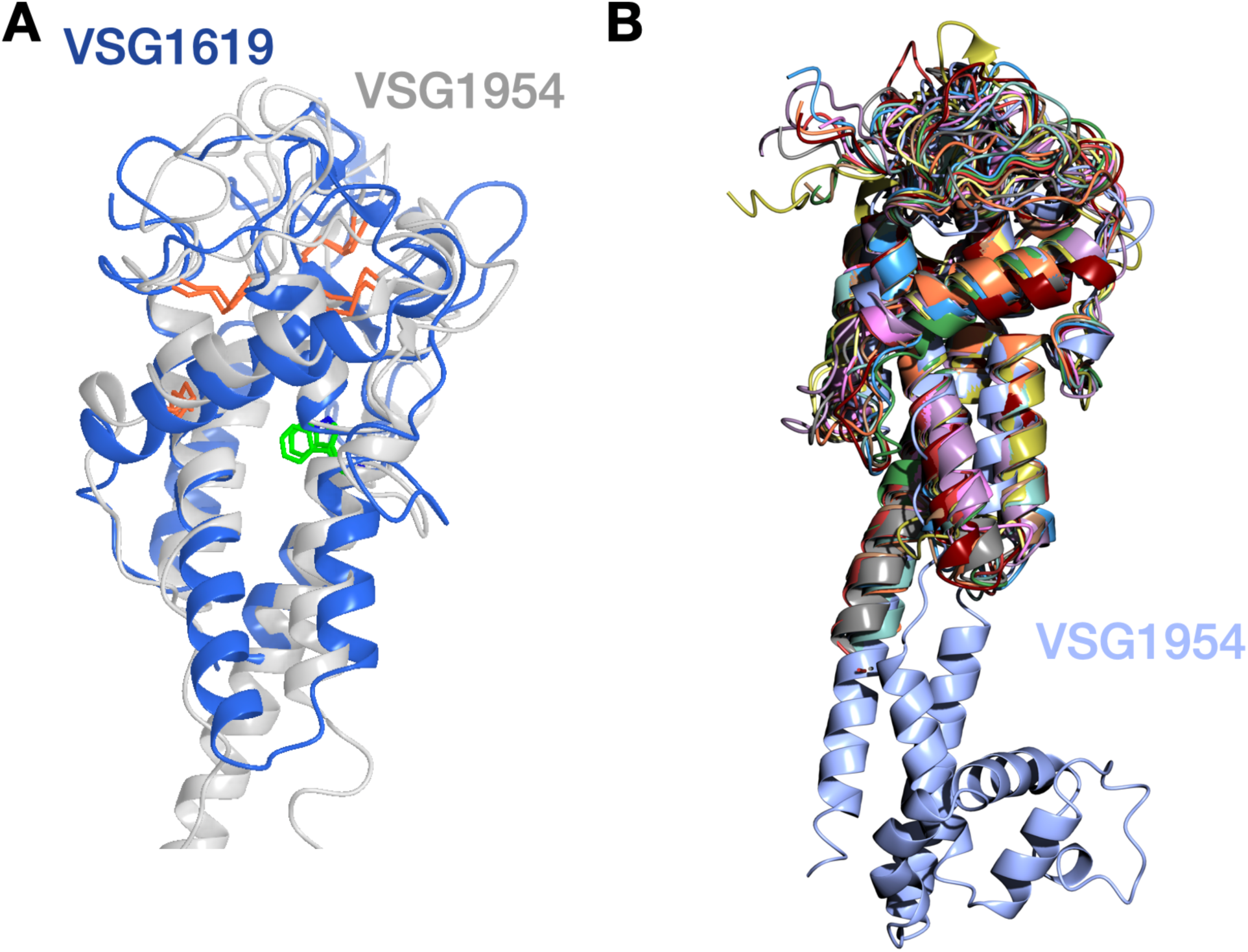
VRP Family Aligned to B-Class VSG. (A) Superposition of VSG1619 (blue) and the top portion of B-class VSG1954 (gray). Disulfide bonds are shown as stick-figures in orange and a conserved aromatic residue in the B-class shown in green. (B) Superposition of 15 VRP proteins (multiple colors) related to VSG1619 and VSG1954 (light blue).

VSG1619 harbors two potential *N*-glycosylation sites per chain (Asn 203 and Asn 233, S5 Fig, [42]), yet no carbohydrate is present in the crystal structure, despite extensive ordered solvent. It should be noted that the location of these possible *N*-linked glycan sites is in what would normally be called the “upper lobe” of the protein, a location that to date has not yielded any *N*-linked glycans in the A or B-class VSGs characterized. It is unclear whether these glycosylations are absent in this protein natively, or lost in the HEK expression system and/or during purification.

### Heterologous Expression of Truncated VSG3327

The HEK expression system was also successful in producing soluble and stable protein of VSG3327, another shortened VSG in the same family as VSG1619 that could not be produced in *T. brucei* VSG expression sites (S2 Fig, S3 Fig panel A and S4 Fig panel A). VSG3327 itself is located on BES 11 between other ESAGs (*Expression Site Associated Gene*s) and described as a VSG-related gene [43]. Interestingly, like VSG1619, it also possesses several potential *N*-linked glycans in positions modeled to be in the “upper lobe” portion of the protein (S5 Fig, A and B). However, in this case the glycans were demonstrably present in HEK heterologously produced protein. The first clue came in the purification as the VRP presented with two distinct bands on SDS-PAGE (S5 Fig, panel C). Mutation of the putative carbohydrate attachment sites (N50A, N192A) reduced the apparent molecular weight on an SDS-PAGE gel, indicating a likely loss of a glycan, and PNGase treatment also showed a decrease in molecular weight (S5 Fig, C and D). Efforts to determine the structure of this VSG are ongoing.

### Expression of a VSG from *T. vivax*

Finally, we have used the HEK system to successfully produce the NTDs of VSGs from the distally related *T. vivax*. Automated predictions of *T. vivax* VSGs often show low confidence (S6 Fig), possibly due to the absence of related genes and structures that can be used for the multiple sequence alignment and starting template in the AlphaFold predictions. For these reasons, we sought to express *T. vivax* VSGs in *T. brucei*, but as with the truncated *T. brucei* VSGs, we were unable to obtain clones for most *T. vivax* genes. An exception was F9WSQ1, which could be expressed in *T. brucei*, but no crystals were obtained with such material to date. However, expression of several *T. vivax* VSGs was achieved with the heterologous system. We present here the structure of one of these, F9WSW0. For the determination of the F9WSW0 NTD, AlphaFold prediction and SignalP were used, indicating that the mature protein starts at Met137 (thus the first 136 residues from the sequence of UniProtKB Accession F9WSW0 were removed and we begin counting the protein here from that residue).

F9WSW0 was solved to 1.74 Å resolution (Methods, S2 Table) and evinces the canonical dimeric arrangement of the *T. brucei* Class-A VSGs in which two monomers that consist of three-helix bundle cores interact along the bundle axis (Figure 4A). The top lobe of F9WSW0 is reminiscent of this class as well, composed of a limited number of short beta-sheets connected by extended loops. However, the top lobe in F9WSW0 is considerably larger than that in VSG2 and related A2-class VSGs from *T. brucei*. While both NTDs are around 100 Å in the long-axis parallel to the bundle, the top lobe in VSG2 is roughly 40 x 50 Å, whereas that of F9WSW0 is 50 x 50 Å, giving the *T. vivax* VSG a larger, more symmetric “mushroom” shape as compared to the more ellipsoidal top lobe of VSG2 (Figure 4B). F9WSW0 is also thicker than VSG2 below the top lobe along the three-helix bundle, with two helices per monomer extending along the same axis as the bundle, widening the “stalk” of the VSG fold. The NTDs of these two domains are roughly the same in sequence length (F9WSW0 with 332 residues and VSG2 with 358). The “additional” sequence in VSG2 that is not seen in the upper lobe and stalk relative to F9WSW0 occurs in the lower lobe of VSG2 with a 100-residue subdomain common to all the Class A2 VSGs from *T. brucei*. F9WSW0, in contrast, does not have this bottom lobe in the crystal structure or in the AlphaFold3 models of the sequence. Instead, the remaining 30 amino acids at the C-terminus of the NTD (missing in the experimental structure) are predicted by AlphaFold 3 to form an extended helix running below the NTD core fold (S6 Fig). Thus, the *T. vivax* protein, while possessing many overall features common to *T. brucei* Class-A2 VSGs, differs from them in being far more “top heavy” with an expanded upper lobe and helical stalk while lacking completely a lower lobe subdomain.

**Fig. 4.**
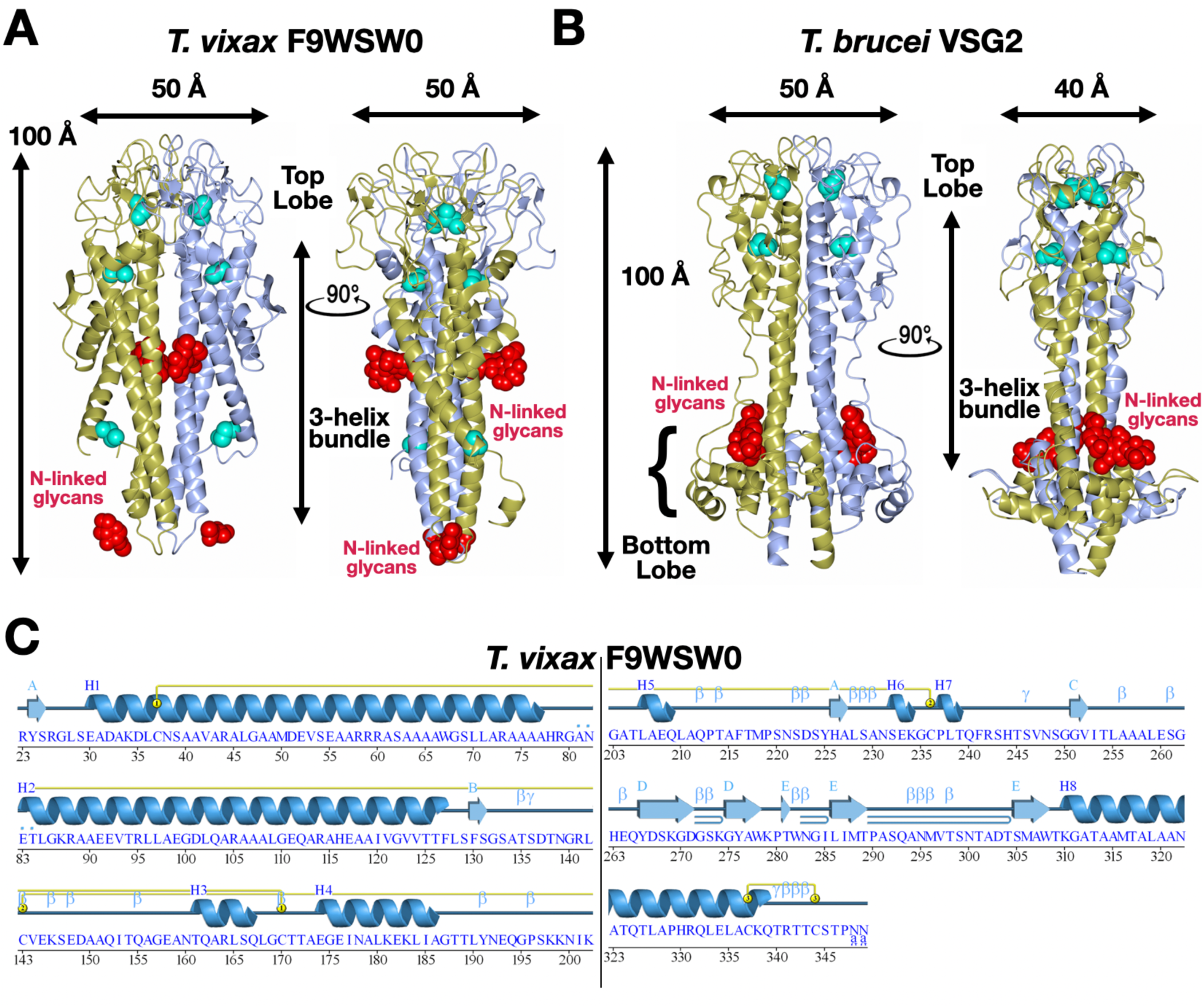
Structure of VSG-F9WSW0 and Comparison to VSG2. (A) Ribbon diagram of VSG-F9WSW0, the two chains of the dimer shown in gold and light blue. *N*-linked glycans are shown in red in space-filling representation. Disulfides are shown in cyan. (B) Ribbon diagram of VSG2, the two chains of the dimer shown in gold and light blue. *N*-linked glycans are shown in red in space-filling representation. Disulfides are shown in cyan. (C) Secondary structure of VSG-F9WSW0 shown above the corresponding sequence, the first part of the protein shown on the left and divided by a horizontal line from the second portion on the right.

Similarly, other structural elements, such as disulfide bonds and *N*-linked glycosylation at the bottom of the NTD, have close parallels between F9WSW0 and *T. brucei* Class A2 VSGs (Figure 4, A and B). Differences arise in that F9WSW0 also possesses a disulfide toward the lower region of the NTD, not seen in any published Class A2 *T. brucei* VSGs. Additionally, F9WSW0 possesses a second *N*-linked glycan near the middle of the NTD, a feature not seen in other *T. brucei* Class A2 VSGs (although similarly placed glycans are present in VSGs of the distinct A1 class) which are to date characterized by only one *N*-linked glycan that is located in the bottom lobe (or, in the case of ILTat 1.24, no sugar at all). In keeping with the general divergence seen between VSGs even in the same organism, the molecular surfaces of F9WSW0 and other VSGs are distinct in electrostatic distribution, hydrophobicity, molecular topography, and other features (Figure 5), consistent with the antigenic distinction between these individual coat proteins.

**Fig. 5.**
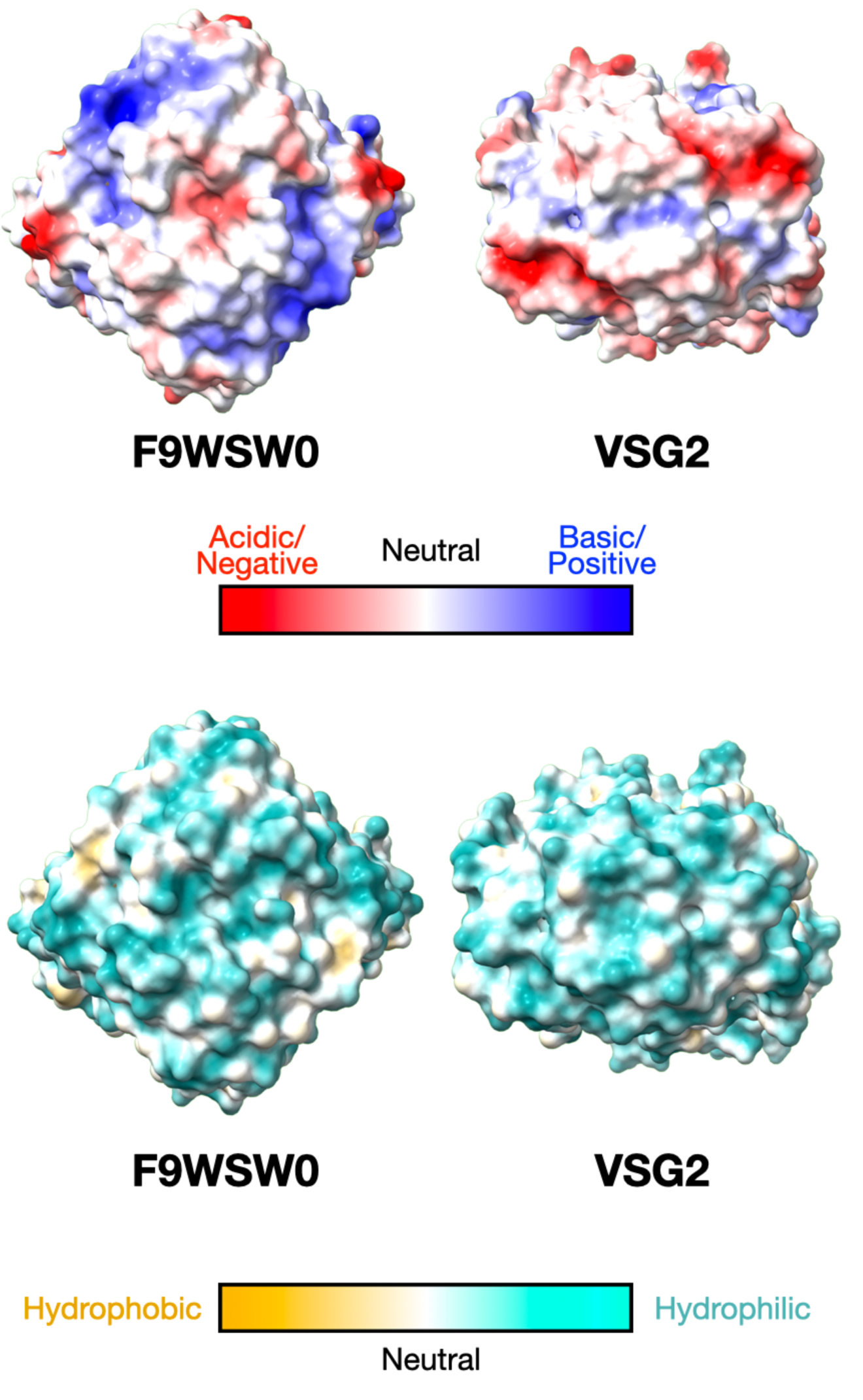
VSG-F9WSW0 and Surface Comparison to VSG2. Molecular surfaces looking down (90-degree rotation from the ribbon diagrams) at the “top” of the VSGs. Electrostatic (coulombic) color is on the left and hydrophobicity (Molecular Lipophilicity Potential, MLP) on the right. ChimeraX was used for these images with identical color mapping for the compared structures.

## DISCUSSION

The central host-pathogen interface in trypanosomiasis is the interaction between host antibodies and the VSG coat of the parasite. Antigenic variation of this coat, obtained through the switching of VSG proteins with distinct antigenic surfaces, is the countermeasure to the adaptive immune response of the host, allowing the pathogen to persist exposed in the bloodstream to the full surveillance and response of defenses to infection. Therefore, understanding the epitope space of the VSGnome is central to developing a robust model of antigenic variation.

That effort has been hindered in part due to the limitations of native cell-culture and recombinant expression systems for VSG proteins beyond *T. brucei*. In some cases, bacterial systems have succeeded for the much smaller VSG CTD [44]. While it has been demonstrated that VSGs can be produced recombinantly in heterologous eukaryotic systems [21–23], it has remained less well-established whether these proteins are largely similar to their natively expressed counterparts in many relevant biochemical aspects. To pursue thorough studies of VSGs across multiple species, and do so beyond the limitations of native expression in the parasite, a more validated, recombinant, heterologous expression system would be very useful.

Our findings here show that using HEK293F suspension cells with standard plasmids commonly used in the production and extracellular secretion of antibodies, VSG NTDs can be produced with high yield and purified to homogeneity. Furthermore, biochemical and structural studies of these recombinant VSG NTDs show them to be very similar to those derived natively from *T. brucei*. This similarity includes the protein fold, disulfide bond location, and *N*-linked glycosylation (confirmed with X-ray structures of HEK-produced VSG2 and VSG3). The one exception was the lack of *O*-linked glycosylation on VSG3, something not unexpected as there appears to be a trypanosome-specific glycosyltransferase for this modification [35], whereas many elements of the core *N*-linked glycosylation pathway are evolutionarily conserved between mammals and trypanosomes [45].

With these “control” VSGs showing the robustness of the heterologous expression system, we then used it to express “problematic” VSGs and VRPs both from *T. brucei* and *T. vivax* that have been challenging to produce natively in *T. brucei*. These led to the determination of structures of a highly truncated VRP (VSG1619) and the *T. vivax* VSG protein F9WSW0.

The crystal structure of VSG1619 revealed it to resemble a truncated VSG coat protein. AlphaFold3 predictions, although many of low confidence, support this architecture for the entire family. Facts such as intact signal peptides and GPI-anchor sequences, absence of apparent deleterious mutations, the expansion into a large family of related proteins with high homology, and presence of several within the BES (yet distal to the telomeric VSG location), are all consistent with a surface-coat role that has diverged from canonical coat VSGs and which has been maintained under selective pressure. Supporting this idea is the observation that we have been unable to substitute canonical VSGs with any of these VRPs to date. Nevertheless, it is possible that these do not have a VSG-like coat function (e.g., antigenic variation) but are VRPs acting as lower-abundance surface-associated molecules, potentially interacting with the host or performing yet uncharacterized functions.

The HEK system shows that it can be used to examine the VSGs from more distally related trypanosomes which lack an effective endogenous or trypanosome-based heterologous expression system (e.g., *T. vivax* VSGs produced in *T. brucei*). The structure of *T. vivax* Fam23 VSG, F9WSW0, reveals many similarities and a few differences to related Class A2 VSGs from *T. brucei*, indicating that these VSGs diverged from a common ancestral family. However, both Fam23 and Fam24 of *T. vivax* show a clear evolutionary connection to the Class A and Class B families of *T. brucei* even at the sequence level. Thus, while structural variation is likely to be seen in the related families, perhaps at times significant, a larger unknown is how the other *T. vivax* family VSGs (Fam25 and Fam26) will assemble in terms of protein architecture and oligomeric state.

Because the HEK system is a robust, widely used platform for recombinant protein expression, we are convinced that it will prove useful broadly across VRP space of trypanosome species. Producing more experimentally-determined VSG structures will also inform machine-learning systems that presently struggle with many VSG proteins from species more distantly related to *T. brucei*, likely due to a lack of training data for these divergent organisms. Providing these structural templates will therefore enhance the use of AI methods in this field and help automate future characterization of the trypanosome coat, the central element in the host-pathogen interaction and source of the unique immune-evasive properties of these parasites.

## Supporting information

VRP Alignments

VRP-family Sequence Analysis

## SUPPLEMENTARY FIGURES AND TABLES

Figures 1-6

Table 1-2

Files ClustalAlignment_VRPsmall.pdf, and VRP_truncated.xlsx

### ACKNOWLEDGEMENTS

Measurements were carried out at beamline 14.1 at the BESSY II electron storage ring operated by the Helmholtz-Zentrum Berlin für Materialien und Energie. We would like to thank the MX team for assistance during the experiments.

## AUTHOR CONTRIBUTIONS

J.P.Z., M.v.S., and C.E.S. conceived, initiated, and coordinated the project. M.v.S. cloned VSG2, VSG3, VSG1619, and VSG3327, conducting the purification, crystallization, and data analysis of VSG2, VSG3 and VSG1619. K.D. purified VSG3327, cloned, and expressed VSG3327 mutants, and M.v.S. and K.D. performed the PNGase experiments. J.B. cloned the *T. vivax* VSGs in *T. brucei*, and produced the F9WSQ1 expressing clone, M.v.S. cloned all the *T. vivax* VSGs in HEK cells, J.P.Z. expressed and purified the protein and crystallized and solved the structure of F9WSW0.

---

**S1 Fig.**
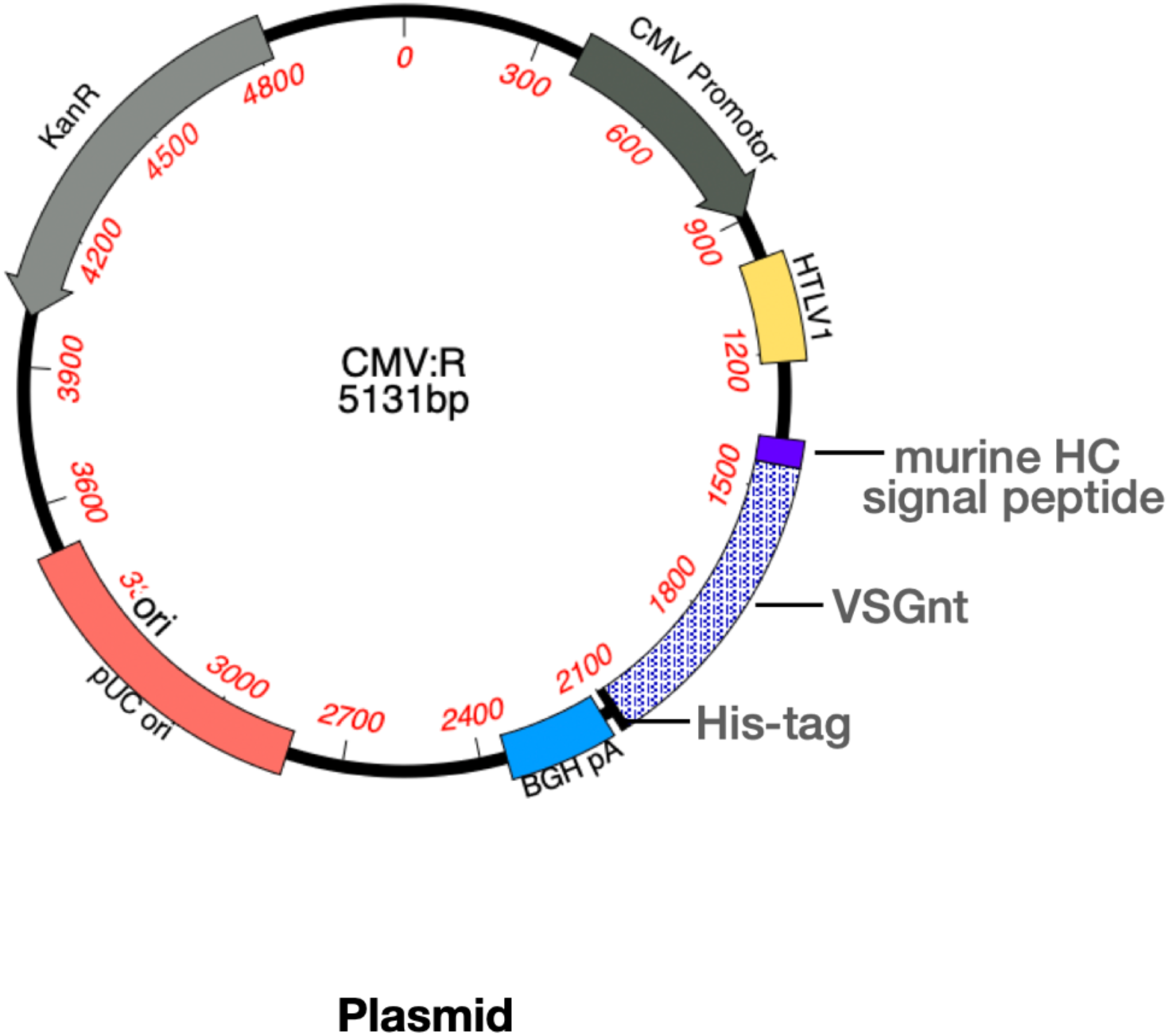
Plasmid map CMV/R for expression of proteins in mammalian cells, derived from VRC01.

**S2 Fig.**
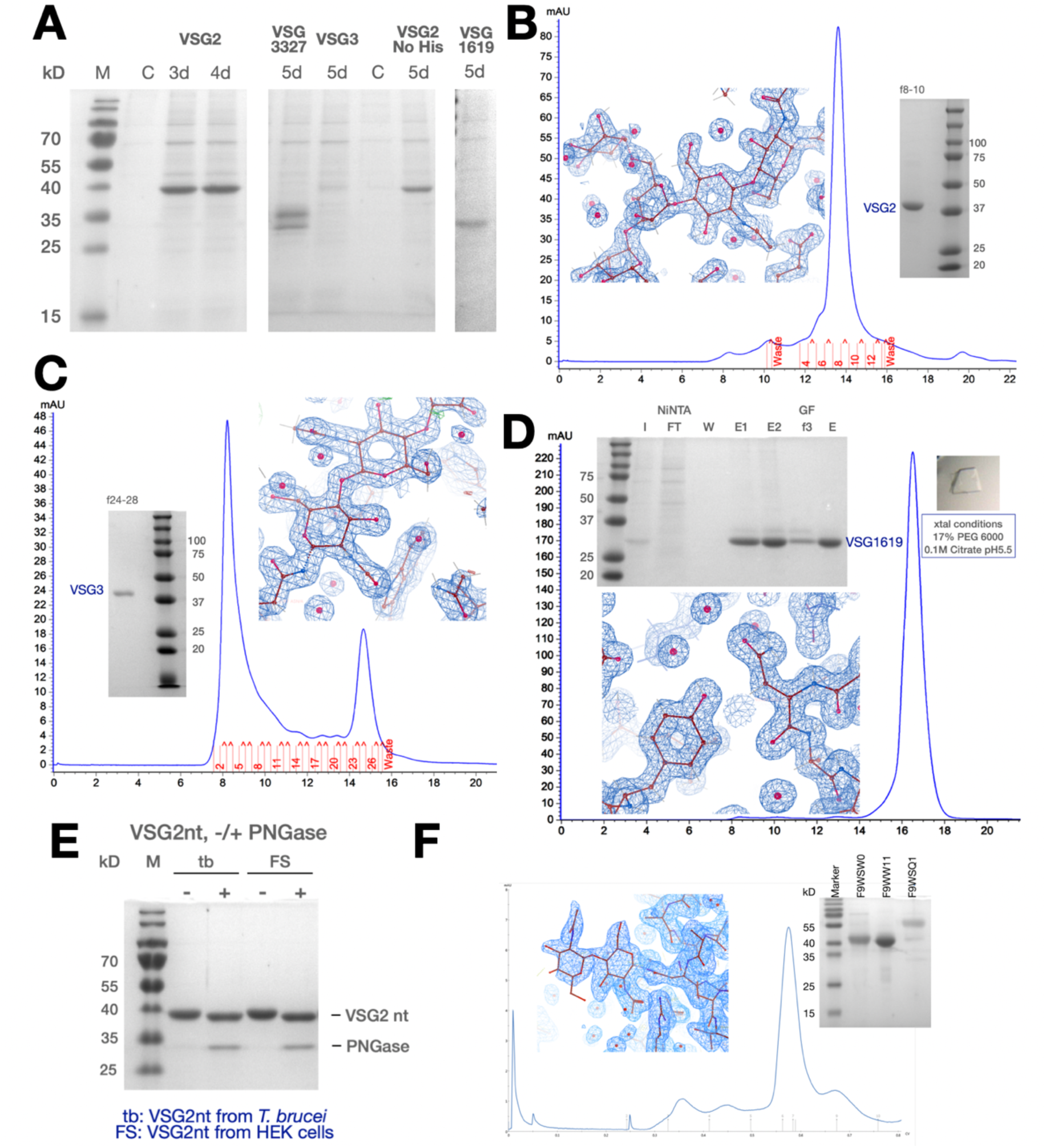
(A) Coomassie stained SDS-PAGE of supernatants from HEK cell expression of different VSGs used for purification of crystallization material (B-D) gel filtration (GF) chromatogram and representative electron density of (B) VSG2 (f8-10 indicated pooled GF fractions), (C) VSG3 (f24-28 indicated pooled GF fractions) and (D) VSG1619, respectively, with a Coomassie-stained SDS-PAGE gel of the final material used for crystallization with the VSG1619 gel also showing samples of the purification steps (FT is flow through, W is wash (Methods), E1, E2 are elution fractions from the affinity purification with the hexahistidine tag (Methods), GF f3 indicates fraction 3 of gel filtration). (E) SDS-PAGE gel of purified VSG2 NTD expressed in *T. brucei* (tb) or HEK293 FreeStyle cells (FS) (lane 2 and 4) and after digestion with PNGase F (lane 3 and 5). (F) F9WSW0 purification (SDS-PAGE Coomassie-stained and Superdex 200 size-exclusion column run) along with representative electron density.

**S3 Fig.**
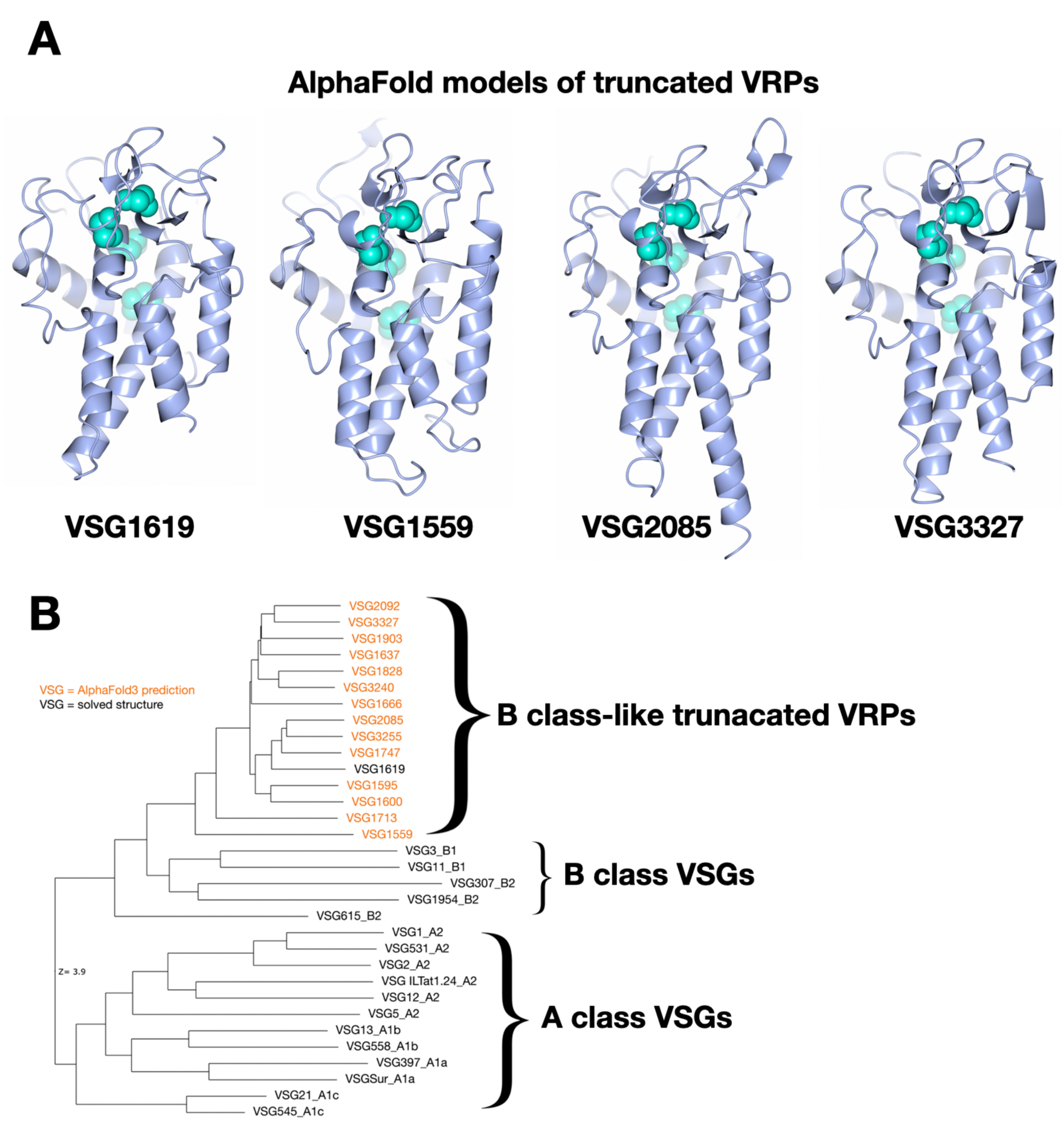
(A) AlphaFold 3 models of four representative VRPs shown as blue ribbon diagrams with predicted disulfides shown in space-filling representation in cyan. (B) Structural similarity dendrogram of multiple VSGs illustrating the B-class relationship of the truncated VRPs examined in this study.

**S4 Fig.**
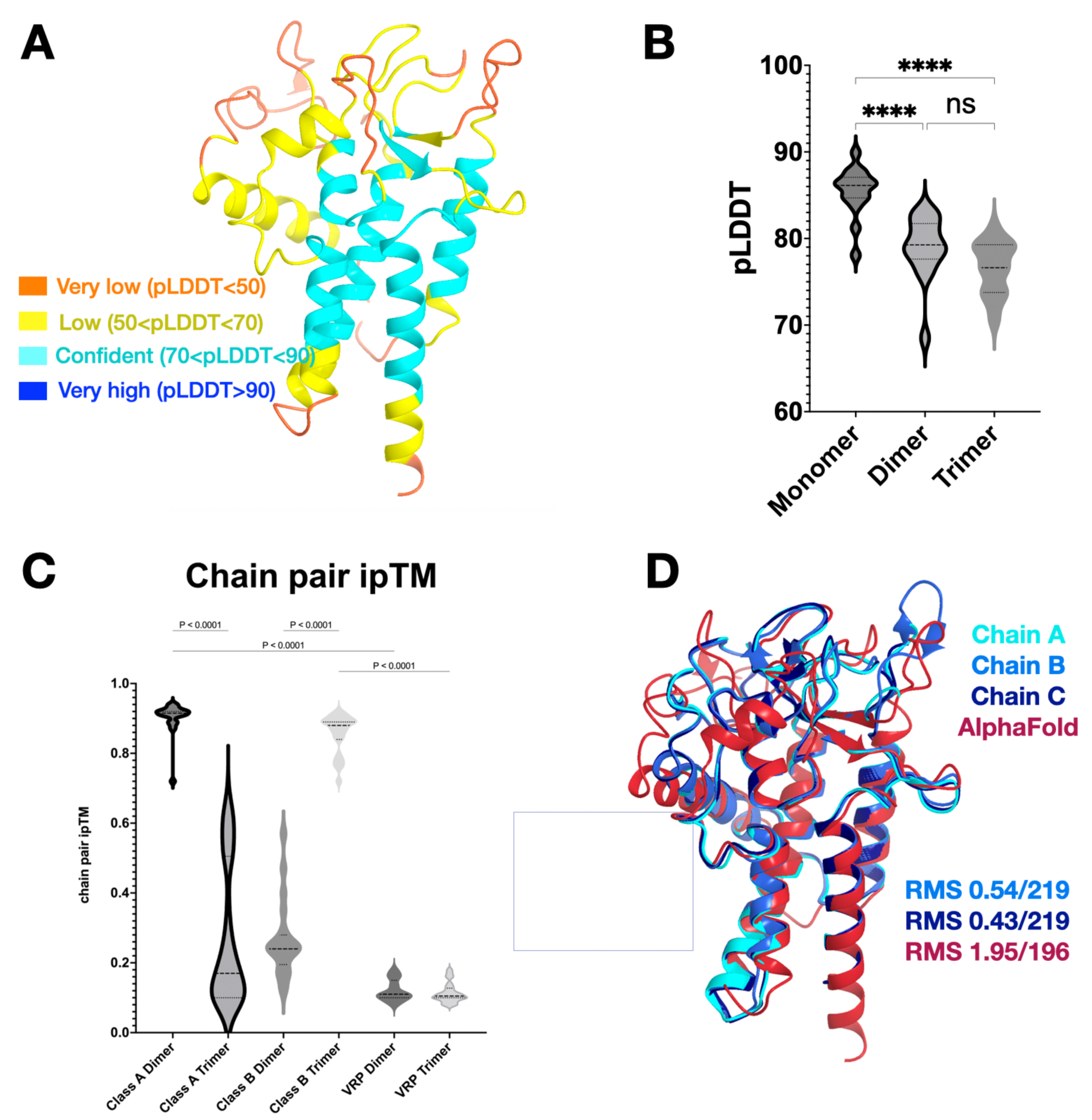
Comparison of predicted structures for small VRPs to the experimental determined structure of VSG1619. (A) VSG1619 AlphaFold3 prediction shown as a ribbon diagram colored by pLDDT value (**p**redicted **L**ocal **D**istance **D**ifference **T**est, a per-residue confidence score (0–100) for the reliability of the structural model). (B,C) Violin plot of pLDDT values and ipTM plot for the homologous family of smaller VRPs studied here comparing AlphaFold3 attempts to produce dimer and trimer oligomeric assemblies (thus, “Class A Dimer” indicates an attempt by AlphaFold3 to produce a dimeric assembly). Comparisons to known dimer and trimer oligomeric states (for Class A and Class B *T. brucei* VSGs, with 21 sequences used for each) are shown in the first four elements on the x-axis, with high ipTM values only occurring for the correct oligomeric state, while the VRP proteins score very poorly for either dimer or trimer associations (with eleven sequences used in the calculation) (D) Structural alignment of the three, independent monomers of VSG1619 in the crystal asymmetric unit and the AlphaFold3 model, showing the root mean square deviation in structure (RMSD) over the Cα atoms of the polypeptide chain.

**S5 Fig.**
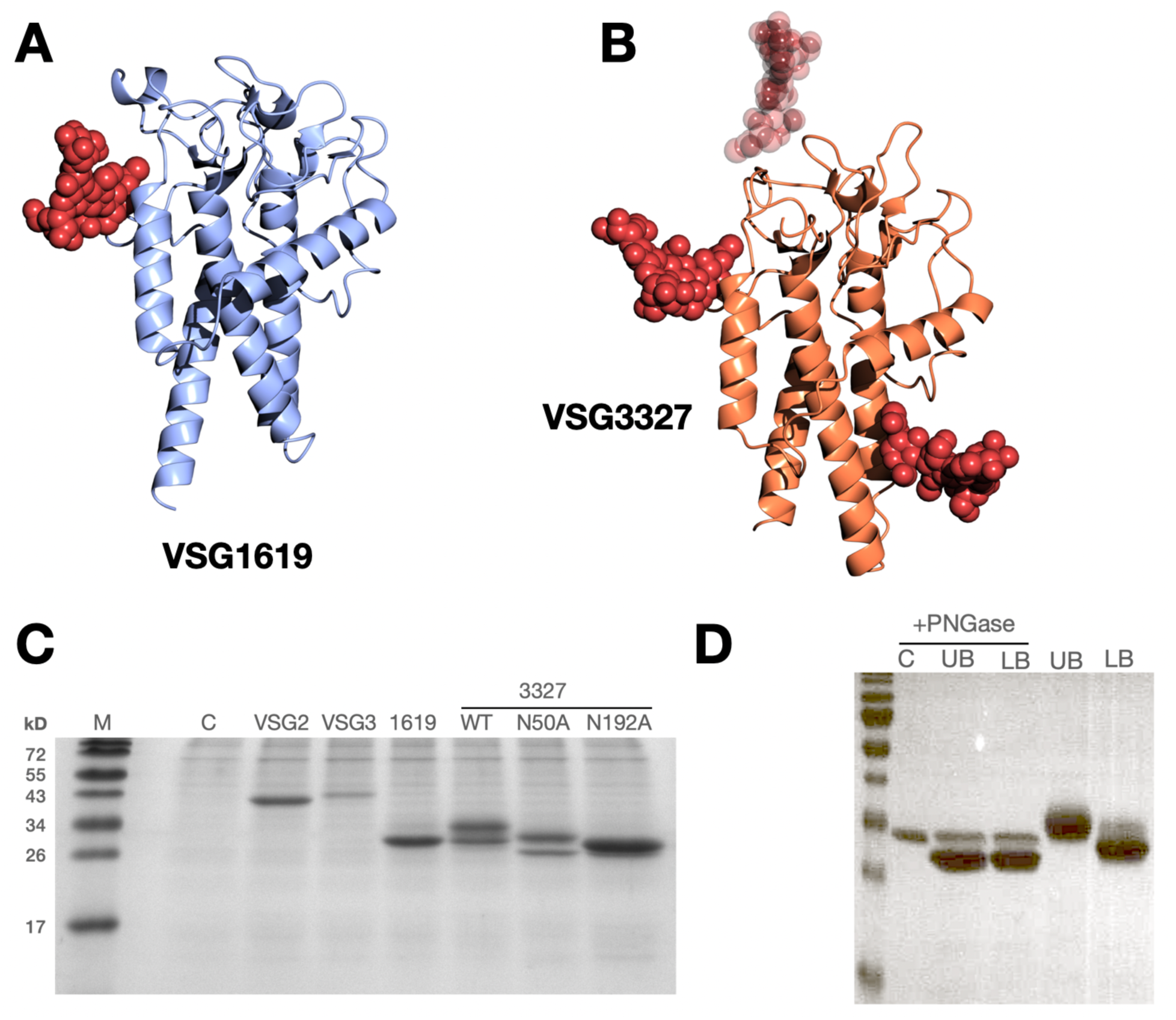
(A) and (B): Predicted locations of N-linked glycans on VSG1619 and VSG3327 (proteins shown as ribbon diagrams) based on NetGly 1.0 [42] (C) SDS-PAGE gel, showing the expression of the N-terminal domains of VSG2, VSG3, VSG1619 and VSG3327 in suspension HEK293 cells (lane 2-5) as well as for two 3327 mutants N50A and N192A both missing a presumed N-linked glycan. Expression levels of the various VRPs vary significantly. The size shift in the N50A sample shows that N50 most likely is glycosylated. The disappearance of other molecular weight bands and the appearance of a single band for N192A indicates that this site is glycosylated. (D) SDS-PAGE gel of purified VSG3327 NTD, upper band (UB) and lower band (LB) (lane 4 and 5) and after digestion with PNGase F (lane 2 and 3). In lane 1 is PNGase only. After PNGase treatment a shift of size is seen for both fractions, indicating that the diversity results from a difference in glycosylation.

**S6 Fig.**
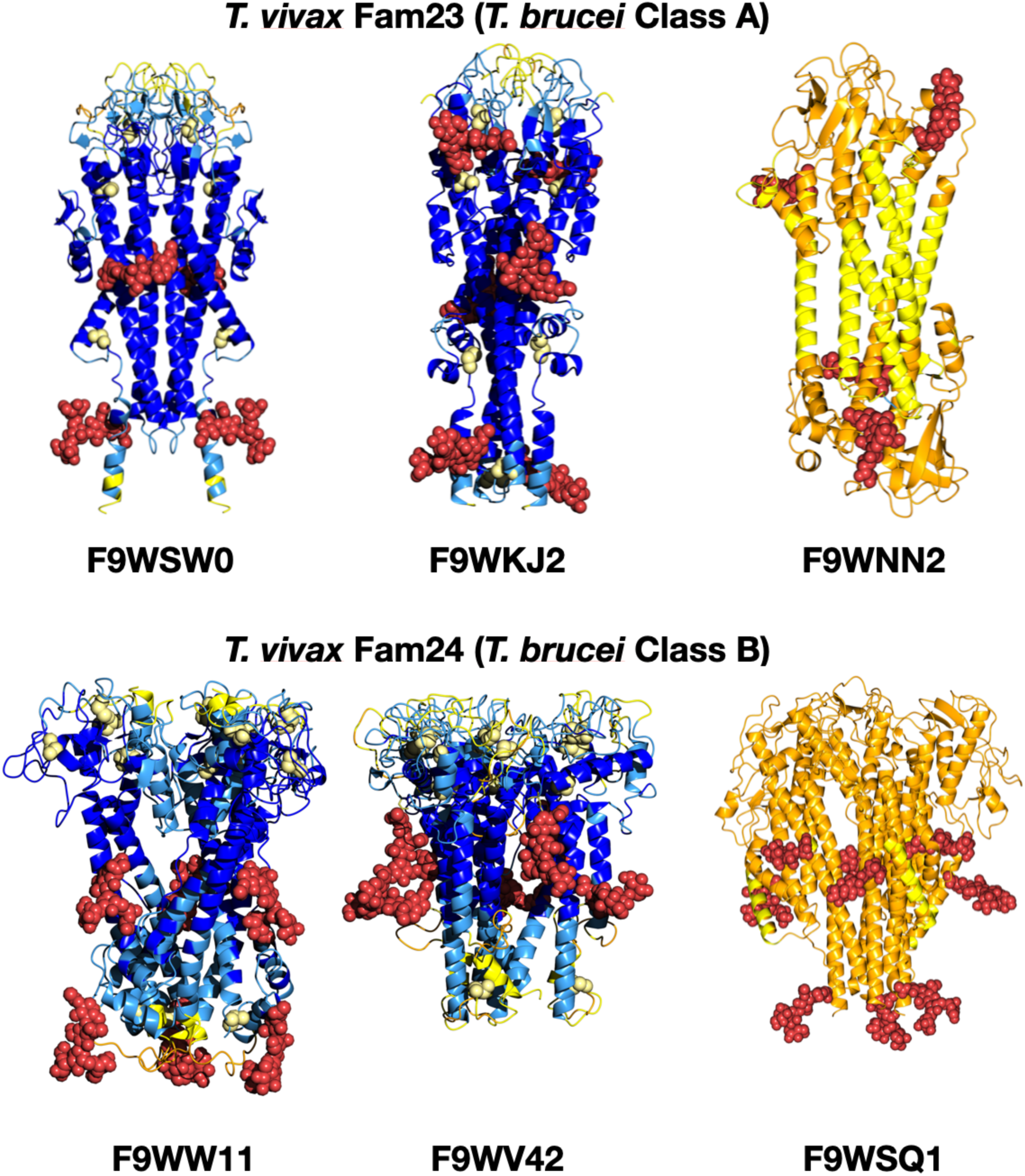
Example AlphaFold3 models of different *T. vivax* VSGs from different families shown as ribbon diagrams colored by pLDDT value as per S4 Fig. Predicted N-linked glycans are shown in red, space-filling representation.

**S1 Table:**
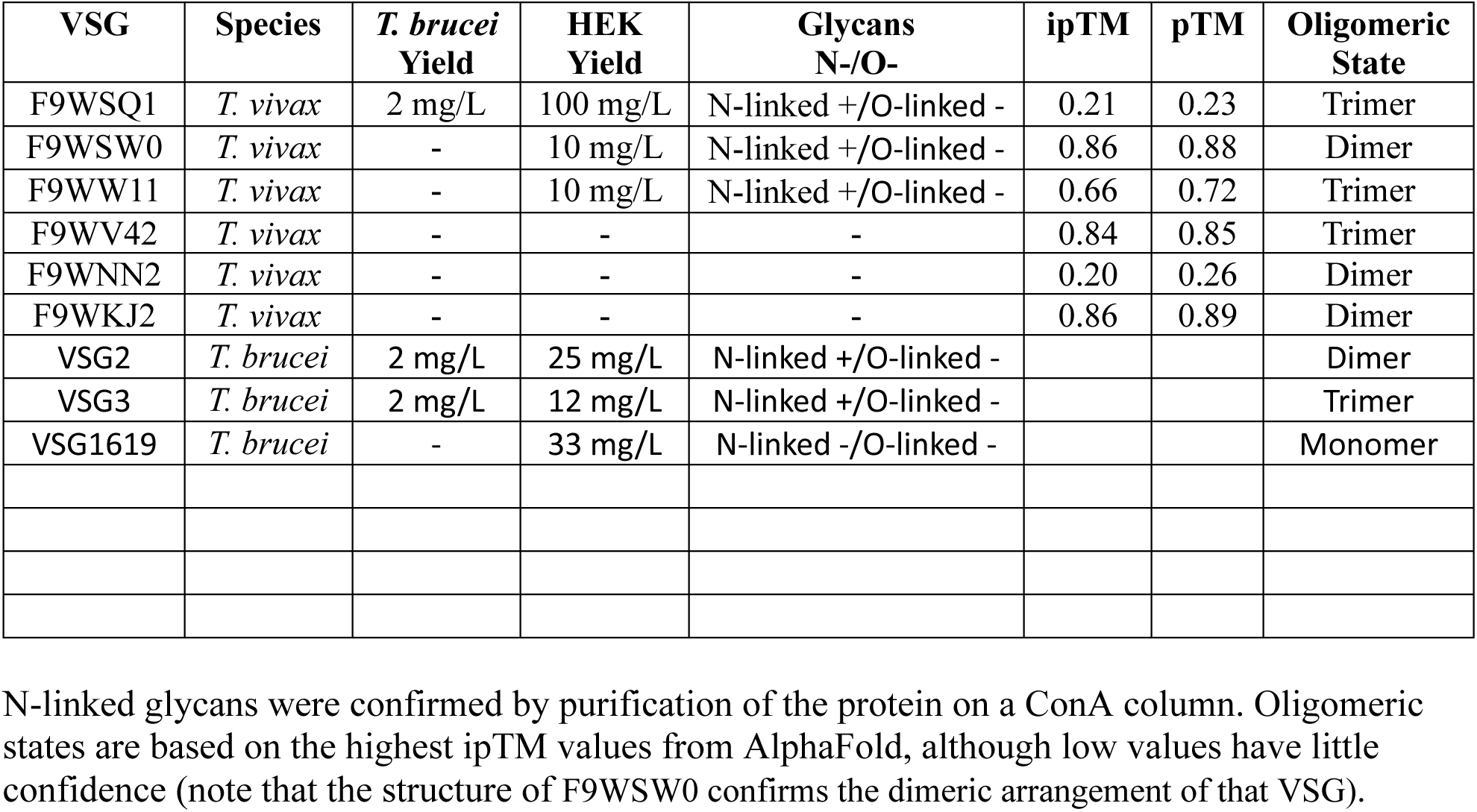
HEK Cell Expression.

**S2 Table:**
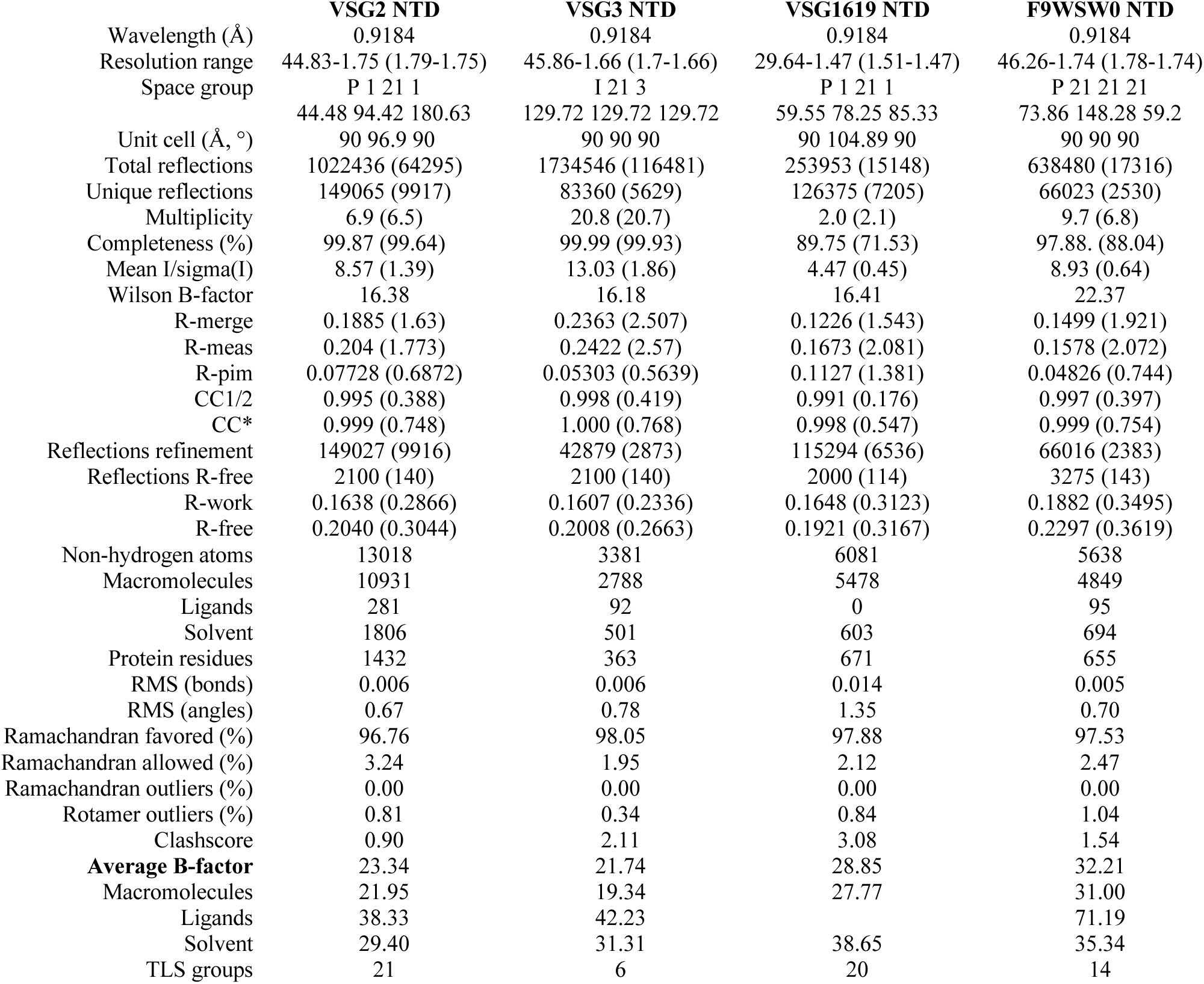
Crystallographic Statistics.

|  | **VSG2 NTD** | **VSG3 NTD** | **VSG1619 NTD** | **F9WSW0 NTD** |
|---|---|---|---|---|
| Wavelength (Å) | 0.9184 | 0.9184 | 0.9184 | 0.9184 |
| Resolution range | 44.83-1.75 (1.79-1.75) | 45.86-1.66 (1.7-1.66) | 29.64-1.47 (1.51-1.47) | 46.26-1.74 (1.78-1.74) |
| Space group | P 1 21 1 | I 21 3 | P 1 21 1 | P 21 21 21 |
| Unit cell (Å, °) | 44.48 94.42 180.63<br>90 96.9 90 | 129.72 129.72 129.72<br>90 90 90 | 59.55 78.25 85.33<br>90 104.89 90 | 73.86 148.28 59.2<br>90 90 90 |
| Total reflections | 1022436 (64295) | 1734546 (116481) | 253953 (15148) | 638480 (17316) |
| Unique reflections | 149065 (9917) | 83360 (5629) | 126375 (7205) | 66023 (2530) |
| Multiplicity | 6.9 (6.5) | 20.8 (20.7) | 2.0 (2.1) | 9.7 (6.8) |
| Completeness (%) | 99.87 (99.64) | 99.99 (99.93) | 89.75 (71.53) | 97.88. (88.04) |
| Mean I/sigma(I) | 8.57 (1.39) | 13.03 (1.86) | 4.47 (0.45) | 8.93 (0.64) |
| Wilson B-factor | 16.38 | 16.18 | 16.41 | 22.37 |
| R-merge | 0.1885 (1.63) | 0.2363 (2.507) | 0.1226 (1.543) | 0.1499 (1.921) |
| R-meas | 0.204 (1.773) | 0.2422 (2.57) | 0.1673 (2.081) | 0.1578 (2.072) |
| R-pim | 0.07728 (0.6872) | 0.05303 (0.5639) | 0.1127 (1.381) | 0.04826 (0.744) |
| CC1/2 | 0.995 (0.388) | 0.998 (0.419) | 0.991 (0.176) | 0.997 (0.397) |
| CC* | 0.999 (0.748) | 1.000 (0.768) | 0.998 (0.547) | 0.999 (0.754) |
| Reflections refinement | 149027 (9916) | 42879 (2873) | 115294 (6536) | 66016 (2383) |
| Reflections R-free | 2100 (140) | 2100 (140) | 2000 (114) | 3275 (143) |
| R-work | 0.1638 (0.2866) | 0.1607 (0.2336) | 0.1648 (0.3123) | 0.1882 (0.3495) |
| R-free | 0.2040 (0.3044) | 0.2008 (0.2663) | 0.1921 (0.3167) | 0.2297 (0.3619) |
| Non-hydrogen atoms | 13018 | 3381 | 6081 | 5638 |
| Macromolecules | 10931 | 2788 | 5478 | 4849 |
| Ligands | 281 | 92 | 0 | 95 |
| Solvent | 1806 | 501 | 603 | 694 |
| Protein residues | 1432 | 363 | 671 | 655 |
| RMS (bonds) | 0.006 | 0.006 | 0.014 | 0.005 |
| RMS (angles) | 0.67 | 0.78 | 1.35 | 0.70 |
| Ramachandran favored (%) | 96.76 | 98.05 | 97.88 | 97.53 |
| Ramachandran allowed (%) | 3.24 | 1.95 | 2.12 | 2.47 |
| Ramachandran outliers (%) | 0.00 | 0.00 | 0.00 | 0.00 |
| Rotamer outliers (%) | 0.81 | 0.34 | 0.84 | 1.04 |
| Clashscore | 0.90 | 2.11 | 3.08 | 1.54 |
| **Average B-factor** | 23.34 | 21.74 | 28.85 | 32.21 |
| Macromolecules | 21.95 | 19.34 | 27.77 | 31.00 |
| Ligands | 38.33 | 42.23 |  | 71.19 |
| Solvent | 29.40 | 31.31 | 38.65 | 35.34 |
| TLS groups | 21 | 6 | 20 | 14 |

