## Supplementary material for "A recombinant expression system for the Variant Surface Glycoproteins enables structural elucidation of challenging classes of trypanosome proteins": VRP Alignments

```
CLUSTAL O(1.2.4) multiple sequence alignment

VSG      Class   Sequence
VSG307   B2      ---------DDENHKEFKQLCRVYNLLTAAVVEQKISSGNSEQRETITAAASRTLANVKK 51
VSG1954  B2      -----ADVAAGSNAESYAVLCTLVQLTKATKPSVPKSKIID---EVYNVAAA------IG 46
VSG1559          -SKAGKKDSQILNENEFKTLCGFVN----------------------------------- 24
VSG1713          LSVPAAETENTLNQREFDVLCSFLN----------------------------------- 25
VSG3327          ------VEEEIVNEGEFKTLCDFVS----------------------------------- 19
VSG2092          -----CVEQEIVNEDEFKTLCEFVS----------------------------------- 20
VSG1828          ------AGGSIVNEEEYKTLCGFVS----------------------------------- 19
VSG3240          ------AGGTIVNEEEYQTLCEFVI----------------------------------- 19
VSG1595          ------VEQDIVNEDEFKTLCEFVG----------------------------------- 19
VSG1600          ------VEQNILNEGEFKTLCGFVS----------------------------------- 19
VSG1666          -------EGSLANEREFATLCGFVS----------------------------------- 18
VSG2085          ------AEQAILNAKEFQTLCGFVG----------------------------------- 19
VSG1619          -------EQAILNAKEFQTLCGFVS----------------------------------- 18
VSG1747          ---------AILNAKEFQTLCGFVS----------------------------------- 16
VSG1637          -------EEGIVNEEEFKTLCEFVS----------------------------------- 18
VSG1903          ------AEQGILNEDEFKILCEFVS----------------------------------- 19
VSG3255          ------AEEGIVNEEEFKTLCEFVS----------------------------------- 19
VSG615   B2      ---------AYENAKQYEALCGAYAITKQAISDAEYIGDTTGD---------PRPKEVED 42
VSG3     B1      --AAADDVNPDDNKEDFAVLCALAALANLQTTVPSI--DTSGL---------AAYDNLQQ 47
VSG11    B1      ----EANIGTGDNVLHRAALCGIIELAGKRAKLETA--LPNFQ---------NELNSILE 45
                             *      **

VSG307           L-NLTAAEEG-KTKVLRDAGQYPTLKKVADDSAAKGYFENVEEAEFQKLREDLEDIENSK 109
VSG1954          L-AIRGDAVV-KNCIDKKDSKYSDLTD--SDIAKKAYTERT----WPVAQAGAAK-LASS 97
VSG1559          ------------------------------------------------------------ 24
VSG1713          ------------------------------------------------------------ 25
VSG3327          ------------------------------------------------------------ 19
VSG2092          ------------------------------------------------------------ 20
VSG1828          ------------------------------------------------------------ 19
VSG3240          ------------------------------------------------------------ 19
VSG1595          ------------------------------------------------------------ 19
VSG1600          ------------------------------------------------------------ 19
VSG1666          ------------------------------------------------------------ 18
VSG2085          ------------------------------------------------------------ 19
VSG1619          ------------------------------------------------------------ 18
VSG1747          ------------------------------------------------------------ 16
VSG1637          ------------------------------------------------------------ 18
VSG1903          ------------------------------------------------------------ 19
VSG3255          ------------------------------------------------------------ 19
VSG615           LYIMTLSDEDYNNKTLTGVTEEGGLEKRKSDILQRRDTY---GREI-------------- 85
VSG3             L-NLSLSSKEWKSLFN-KAADSNGSPKQPPEGFQSDPTW---RKQWPIWVTAAAALKAEN 102
VSG11            L-NMTAAEPTWLDQFR-DKDDRSKPRDLTKQPLPKDTNW---ADHWTAWAKAALPLLNDE 100


VSG307           GTGETFAKTYGTPFSDSQKEAIRAPLTFMAQAATTIHGKLTAVYKQATL-ARQQAQLAFS 168
VSG1954          GEKEKYASWTHRKYTEKQKLKV----HVLTAAISDVKQRADKLNKP--D-KLAELTGALS 150
VSG1559          ----------------------------LTFEIQKLASEGKPTLGG--L-KPESVNESVN 53
VSG1713          ----------------------------LTLRIHEWVEKE-RLTAE--V-AN-DFDSSAM 52
VSG3327          ----------------------------LTQRINESLRGM-EEKSG--V-SVSFLQKKVT 47
VSG2092          ----------------------------LAEQINESLKGI-KEKSG--V-KMASLQKRVT 48
VSG1828          ----------------------------LTQQINSSL-EI-KGKSG--V-SVASLQKKVN 46
VSG3240          ----------------------------LTQQINSSL-EI-KGKSG--V-NVASLQKKVD 46
VSG1595          ----------------------------LTERINELLRGM-EGNSG--V-SVSSLQKKVT 47
VSG1600          ----------------------------LTERINELLRGM-EGNSG--V-SVSSLQKKVT 47
VSG1666          ----------------------------LTHQIRESVYKV-REKSE--V-DVILLQKKVA 46
VSG2085          ----------------------------LTGQINELLERM-KGNSG--V-DVLLLQKKVK 47
VSG1619          ----------------------------LTERISGLL-DM-KGNSG--V-DVLLLQKKVK 45
VSG1747          ----------------------------LTERISGLL-DM-KGNSG--V-DVLLLQKKVK 43
VSG1637          ----------------------------LAERINVSLEGM-RGNPG--V-DATSLQQKVK 46
VSG1903          ----------------------------FVGQISDSLETM-KGKSK--A-DVASVQKRVK 47
VSG3255          ----------------------------LAERINVSLEGM-RGNPG--V-DATSLQQKVK 47
VSG615           ---------HSIPANSEARAAAHVAIKRLFYKAGNLSANIAAAISSIKA-DTRSAGEALN 135
VSG3             KEAAVLARAGLTNAPEELRNRARLALIPLLAQAEQIRDRLSEIQKQNEDTTPTAIAKALN 162
VSG11            THQAKLKEYKLAGLQPEKLERARNTIRRLTAEAVAKAQDPTVAESTADLTTEEDLQKQIN 160
                                             :

VSG307           KAVYGEKVLAGKDAALMKPDKPLPDPTTAANFPWGAADDRDAVCKNPTVNSGKAGSALAI 228
VSG1954          NSLYGNGK----------------SNADTATLPAGGSHISMCGPADGTQGGSIVGKALKF 194
```

```
VSG1559       TILYGNKG-----------------STEIGFE--GENDRKDYCGDR-QNRAHKYAGKSLIK 94
VSG1713       KVLYGSDS-----------------DGSLNWT--GKEKREADCGYT-SIFGGGFGGKALAL 93
VSG3327       DVLFGTGI-----------------GDVSKMK--WQPHRDMDCGSN-TGNRAFWAGEALMK 88
VSG2092       DILYGTGV-----------------SHVDKMV--WKRYRGQDCGED-NRDRPAVAGKALVK 89
VSG1828       DILFGANV-----------------DDVSNMK--WEFYRELDCGKD-SGGRVALAGVTLVK 87
VSG3240       DILFGANV-----------------DDVSKMK--WGVHRELDCGQD-SGGRVALAGETLVK 87
VSG1595       DILYGTGV-----------------SAVNEMR--WNRHREMDCGQD-NRVKPASAGRALVQ 88
VSG1600       DILYGTGV-----------------SAVNEMR--WNRHREMDCGQD-NRVKPASAGRALVQ 88
VSG1666       DILFGTGI-----------------SDVGSMK--WQQYRQQDCGQD-SGGRTFPGGETLVK 87
VSG2085       GILFGTGI-----------------GDVNKMS--WQLYRQLDCGQD-SWPPRTRGGEALAR 88
VSG1619       DILFGTGI-----------------GDVGKMN--WGYYRELDCGQD-SGNHQTKGGEALMK 86
VSG1747       DILFGTGI-----------------GDVGKMN--WGYYREMDCGRD-SGNPRTRGGEALVR 84
VSG1637       SILFGAGI-----------------SDVSHME--WGFHREMDCGRD-SEDRRFLAGETLVK 87
VSG1903       DILFGANV-----------------DDVNKML--WKVHREMDCGQE-SGNQRTHGGKALVR 88
VSG3255       SILFGAGI-----------------SDVSQME--WGFHREMDCGQD-SGGIPASAGRSLVK 88
VSG615        RARCGQADCKAP------------------DQKWFETRSKACSGT----GEQKQGMTIAS 173
VSG3          KAVYGQDKETGAVY----------NSADCFSGNVADSTQNSCKAG----NQASKATTVAA 208
VSG11         QAVYSKDTEPDDDF----------NGYTAFEGKASTNRQTICGSA----VAGSKATNAMD 206
                   .                                   .              .

VSG307        DMVCICTKKQATP-------QQLCGTALTGGSTVIDSSSSQSKAHAAWKALSAACTKMAN 281
VSG1954       DLICLCGKQSAD----SGTGEKACHEFS---PLPATAIAENAAINADWATIEQGCKTVAG 247
VSG1559       DILCLCKPHRRQGQNGAGA-EDLCFSGNKW-YKGGEVWSNKEQAKQHWEKIRLHCSQLPK 152
VSG1713       DLLCLCRPSGNG----DHQ-KNLCCEANAW-YDPAESWGNKDRAEESWGKIKQKCTEQIN 147
VSG3327       DLLCLCERTDGL---HSSP-EHLCYRGRTK-RHNADNWQSPTNRKNTWNELQNKCKKGGI 143
VSG2092       DLLCLCEGRDGR----STL-KDICYTGNDK-RHNSAKWEDSVHHKNTWNELQNKCERGRS 143
VSG1828       DLLCLCEGRDGT----SPP-GDLCYTGNNK-RYNSQTWGSSTNHKNTWNELQNKCASGRS 141
VSG3240       DLLCLCEGRDGT----SPP-GDLCYTGNNK-RYNTETWRSSTNHKNTWSELQNKCASGRS 141
VSG1595       DIICLCDGRDQK----PSK-KDLCYKESGE-HRRYTYWWDSQGHGNTWSHLRNGCKTERG 142
VSG1600       DIICLCDGRDQK----PSK-KDLCYKESGE-HRRYTYWWDSQGHGNTWSHLRNGCKTERG 142
VSG1666       DLICLCDGTDLQ----PRR-DALCYAENTE-QSRYTGWKNTLNHEKTWSRLQGKCEADHR 141
VSG2085       DIICLCEGTDQQ----PNL-RDLCYAGNIK-QYDFNTWGGSENHRNTWSSLKSECAKEHG 142
VSG1619       DLVCLCEGTDQQ----SPP-GGLCYEGNTK-KNSRGGWNNAEAHRNTWSSLKSECTREHG 140
VSG1747       DLICLCEGTDRQ----SPP-RGLCYEGNTR-TVSPGKWDNAEAHRNTWSSLKSECTRERG 138
VSG1637       DLVCLCEVTGLQ----RPG-KDLCYAGNAR-KFSLQEWPTVQKHRSTWDDSRNKCKMRHT 141
VSG1903       DLICLCEGTNRQ----PNL-KDLCYTGNAR-KFSSQEWPTTQKHRSTWDDLRSRCITGSG 142
VSG3255       DIICLCEGTGQQ----PNL-RDLCYTGNVR-KVNSQEWPTTQKHRSTWDDLRSRCITGSG 142
VSG615        DISCLCSAATGE----------TLCSAAATGG-TYRGGEGTAANAQTDWSTTIADCDRNVE 223
VSG3          TIVCVCHKKN-GG----NDAANACGRLIN---HQSDAGANLATASSDFGDIIATCAARPP 260
VSG11         ALFCVCADDRTNG----ADAGKACVAGTAPG-TGWNPGVTATPTGTMLQKVRKLCNTHGK 261
                 : *:*                *                                  *

VSG307        EATGGDQKMQLTTELASVEAMRGQNTIVITGSPGYQALAAKTHNFFGAFVVATTTASDCD 341
VSG1954       APSLTPESIH-----AALQAFYR-------HAG-VPK-GNTRNRYTTVGAPAGSGATGCD 293
VSG1559       RQQ----HIQ-----NQLYHLKEKVQAVLRTATEKK--GRESDQNVRLGGHQATEVKQCS 201
VSG1713       IESIGFESLT-----TTVQNLRKKLNEMTEKT--------TERETKRLGHGYEPITKSCG 194
VSG3327       EGA----PTR-----TEFQDMRKQLEAGIKKRNDSQ-----HGEYCIYGGEKAKTLRICD 189
VSG2092       EGV----PTR-----TEFQGMRKRLETRIKERKRSG-----GRGNYIYGGNKDDAPTLCD 189
VSG1828       GVP----PTR-----TEFQGMRITFETGIKERKRSG-----GHGYYTYGGNKDKVLSVCD 187
VSG3240       GVP----LTR-----TELQGMRITFETGIKERKRSG-----GGGYYTYGGNKDKVLSVCD 187
VSG1595       GSA----PKE-----AVFKELKRQLQAGLKTYP-----GGNESAYYTYGGHTNSGLQTCN 188
VSG1600       GSA----PKE-----AVFKELKRQLQAGLKTYP-----GATESAYYTYGGHMNSGLHTCR 188
VSG1666       EGT----PTR-----AEFENKKEQLRSGIKGRKDKE------GTYYTYGGNAKNGLQICN 186
VSG2085       EGL----LAE-----SELQEMKKQLKERLKERKDMRT-SAREGTFYTYGGGKEYGLHVCD 192
VSG1619       EGL----LAE-----SEFQERKKQLKERLKERKDMRA-GTREGTFYTYGGNTSNGLQTCN 190
VSG1747       ESL----PIE-----IEFRKKKEKLKERIRDRADRSQ-G---EHLYTYGGGKEYGLQTCN 185
VSG1637       EGL----PTQ-----TEFQEKRKRFETRIKGCQSSS----GR-EYYTYGGDKTKTLSVCD 187
VSG1903       KGV----PSE-----TEFHENKVQFRMRIKKRKNSD----GREHLYTYGGGKEYGLHTCN 189
VSG3255       KGV----PSE-----TEFHENKVQFRMRIKKRKNSD----GRQHFYTYGGGTNNGLQTCN 189
VSG615        GKAPSPAAIE--AAIAVFRAALGNAEFT-------------KANSRKAFVLGHGSASDCN 268
VSG3          -KPLTAAYLD--SALAAVSARI---R-----------------FKNGNGYLGKFKATGCT 297
VSG11         -TTLSAAAIE--GRLTAVGNLL---T-----------------RGSATSILGSFLATDCS 298
                                .                                        *

VSG307        TDDANVVNTGGKGPCIGYSAY--L--KTAAGIPWINHAKTGHSKLQEADDLF-------K 390
VSG1954       GIG-----GSNGGKCAAYNKAQFE--A-GTGPYWATQMKAAAETLVELRGQEQKLAALEA 345
VSG1559       ATT-----GKGNSVCLMYNITSGT--LTNHSIPWLKHLEELAEKIPNDTRTN-------- 246
VSG1713       GSG-----SANNAPCVFYKMEGRG--ANQVKIEWLTQLEDLVKRLKT------------- 234
VSG3327       GRR-----TETNGICVLYPRGPDG--DNASGIEWLNKLERLVKNVEKINQ---------- 232
VSG2092       GQE-----SQTNGICVMYPEGSEE--DSTIGIKWLGELKDLVEEVEEMSKSASTHTE--- 239
VSG1828       GQE-----SETNGICVMYPEGSDK--ESTIGIKWLSELEDLVERVGEMRKNANADTDTAA 240
```

```
VSG3240         GHQ-----TETNGICVLYPRVSXQ--DNTRGIKWLSELEGLVERVEEMSKGANADTE--- 237
VSG1595         GFT-----TSTDGICVMYQRGSDQ--DSTIGIKWLNNLEDLVKKVEEMKTASTRAE---- 237
VSG1600         GTP-----TETDGICLLYRRVSDG--DNASGIEWVGKLEDLVKQVEEISRSANTGADTN- 240
VSG1666         GIA----TTEADGICVLYKRGSDG--DNTSGIEWLNKLEELVKIVEGMDQSTNSD----- 235
VSG2085         GYP-----TDSNGICVMYPKGSDG--DNASGIEWLDKLEDLVKEVEKMNKDESAGGNTK- 244
VSG1619         GAE-----SENDGICVLYPRGSNE--DNASGIEWLNKLEDLVREVEEMSKDESKG----- 238
VSG1747         GAE-----SDNDGICVMYPRGSHG--DNASGIEWLKKLEDLVKEVEKMNKDESAGGN--- 235
VSG1637         GQE-----SQTNGICVMYPKLPNQ--GSAGGIKWLTNLEDLVKTVEKMNKDETASRRTE- 239
VSG1903         GAE-----SENDGICVLYPRGSNE--DNASGIEWLNKLEDLVKEVEEMSKDESTSGSTK- 241
VSG3255         GAK-----SENDGICVMYPKGSDG--DNTSGIEWLKKLEDLVKEVEKMNKDESAGGNTK- 241
VSG615          GGT-------SSAACVDYTNKLAR--GTINDIPWIEQLRTAAAKLAGVAGTRAQLD---- 315
VSG3            GSA-------SEGLCVEYTALTAATMQNFYKIPWVKEISNVAEALKRTEKDAAEST---- 346
VSG11           GDQ-------GSGMCVAYTEVTDAKGTPTKDIPWMQKLDSVRIKLQKHERAVEKLG---- 347
                            . *  *                *  .       :

VSG307          ESLVLLTEVQNIEKQMQSLLLMSKSLTQATQ-A--------------------------- 422
VSG1954         EALALNSTLDGMQHD--------------------------------------------- 360
VSG1559         ------------------------------------------------------------ 246
VSG1713         ------------------------------------------------------------ 234
VSG3327         ------------------------------------------------------------ 232
VSG2092         ------------------------------------------------------------ 239
VSG1828         ------------------------------------------------------------ 240
VSG3240         ------------------------------------------------------------ 237
VSG1595         ------------------------------------------------------------ 237
VSG1600         ------------------------------------------------------------ 240
VSG1666         ------------------------------------------------------------ 235
VSG2085         ------------------------------------------------------------ 244
VSG1619         ------------------------------------------------------------ 238
VSG1747         ------------------------------------------------------------ 235
VSG1637         ------------------------------------------------------------ 239
VSG1903         ------------------------------------------------------------ 241
VSG3255         ------------------------------------------------------------ 241
VSG615          ----------GMRQE----MRIIEDQAWQAF-ALATI----------------------- 337
VSG3            ----------L----LSTWLKASENQGNSVAQKLIKVGDSKAV----------------- 375
VSG11           ----------KPQHDLKTILTLAKD---PAYLQLASVGTRHLETTKQRVSNEQGKTQQTQ 394


VSG307          --     422
VSG1954         --     360
VSG1559         --     246
VSG1713         --     234
VSG3327         --     232
VSG2092         --     239
VSG1828         --     240
VSG3240         --     237
VSG1595         --     237
VSG1600         --     240
VSG1666         --     235
VSG2085         --     244
VSG1619         --     238
VSG1747         --     235
VSG1637         --     239
VSG1903         --     241
VSG3255         --     241
VSG615          --     337
VSG3            --     375
VSG11           QT     396
```
